# Topoisomerase II inhibitors CX-5461 and Doxorubicin differ in their cardiotoxicity profiles

**DOI:** 10.1101/2025.07.20.665626

**Authors:** Sayan Paul, José A. Gutiérrez, Alyssa R. Bogar, Michelle C. Ward

**Affiliations:** Department of Biochemistry and Molecular Biology, University of Texas Medical Branch, Galveston, Texas, 77555, USA

**Keywords:** cardiovascular disease, CX-5461, Doxorubicin, cardiotoxicity, topoisomerase II, global gene expression

## Abstract

CX-5461 (CX) is a chemotherapeutic drug currently under investigation for the treatment of late-stage cancers. While CX was first described as an RNA polymerase I inhibitor, it has recently been shown to primarily inhibit the beta isoform of topoisomerase II. This isoform is also inhibited by anthracycline drugs including Doxorubicin (DOX) and mediates the toxic effects of these drugs on the heart. It is unclear whether CX will similarly cause cardiotoxicity. We therefore designed a study to test the effects of CX compared to DOX on iPSC-derived cardiomyocytes from six individuals. While both CX and DOX induce cell death in cardiomyocytes, CX is 20-fold less cytotoxic than DOX. At sub-lethal doses, DOX induces DNA damage, while CX does not. Transcriptome profiling following treatment with two sub-micromolar concentrations of both drugs over time reveals that DOX induces thousands of gene expression changes compared to hundreds induced by CX. Comparison of gene expression trajectories across drugs reveals that genes that respond to CX also respond to DOX, while most DOX response genes are drug specific. Shared response genes correspond to pathways related to chromosome segregation and DNA replication. CX does not affect the expression of any of the genes in functionally-validated loci associated with DOX-induced cardiotoxicity. Our data demonstrate that CX treatment of cardiomyocytes induces gene expression changes that mirror a subset of those induced by DOX; however these changes do not coincide with the cardiotoxicity observed with DOX treatment.

## Introduction

CX-5461 (CX) is a compound that was first described as a selective inhibitor of RNA polymerase I (1). RNA polymerase I is involved in transcribing ribosomal DNA genes, which are highly transcribed in cancer cells. CX has therefore been established as a therapeutic agent to selectively target cancer cells including those from B-lymphoma (2), acute lymphoblastic leukemia (3) and colorectal cancer (4). Given the *in vitro* efficacy of CX, it is currently in clinical trials for use in advanced solid tumors with DNA repair deficiencies (5), and advanced hematologic cancers (6). Phase I data from both trials suggests that CX is safe for further consideration.

Following the initial description of CX as an RNA polymerase I inhibitor, CX has been implicated in other cellular processes including activation of the DNA damage response. In ovarian cancer cells it activates replication stress and the DNA damage response (7), and these effects are enhanced when used in combination with a topoisomerase I inhibitor that initiates single-strand DNA breaks (8). Similarly, CX enhances radiation sensitivity of cancer cells through effects on DNA damage and not RNA polymerase I inhibition (9). The DNA damage-associated effects of CX are not restricted to cancer cells as CX treatment of vascular smooth muscle cells activates the homologous recombination DNA damage response pathway (10). Mechanisms behind the toxicity induced by CX in DNA repair-deficient tumors include action as a G-quadruplex ligand resulting in stabilization of G-quadruplex DNA structures (11).

It has recently been demonstrated that the primary mechanism of CX toxicity is through topoisomerase II (TOP2) poisoning and not RNA polymerase I inhibition (12–14). This is perhaps not surprising given that CX was originally derived from fluoroquinolones that interact with TOP2 and G-quadruplexes (15). There are two isoforms of TOP2 – TOP2A and TOP2B. CX preferentially targets the TOP2B isoform (14). TOP2A has also been suggested to mediate effects of G-quadruplex ligands including CX (16), and the DNA-damaging effects of CX in ribosomal regions (17).

TOP2 inhibitors, including Doxorubicin (DOX), are effective anti-cancer drugs; however, they can cause off-target effects on the heart given the high levels of TOP2B expression in this tissue (18). Because CX targets TOP2B, it has been suggested that CX might cause adverse effects on the heart (14). CX treatment also induces mutations across human cell lines further questioning the safety profile of the drug (19).

The risk of cardiotoxicity is an important consideration for any drug in development. DOX-induced cardiotoxicity is a well-established clinical phenomenon that can be effectively modeled *in vitro* using induced pluripotent stem cell-derived cardiomyocytes (iPSC-CMs) (20). The role of TOP2B in mediating the cardiotoxicity associated with DOX has also been demonstrated in this system (21). We have previously shown widespread global transcriptional effects of sub-micromolar treatment of TOP2 inhibitors including DOX and structurally-related anthracyclines epirubicin and daunorubicin, and the anthracenedione, mitoxantrone, on iPSC-CMs (22). It is unclear whether CX, which similarly inhibits TOP2B, will induce similar effects on the transcriptome of iPSC-CMs.

We therefore designed a study to determine the effects of CX treatment on iPSC-CMs from six healthy individuals compared to DOX. We found that while CX is less cytotoxic than DOX, and induces an order of magnitude fewer gene expression changes, CX response genes are shared with DOX.

## Materials and Methods

### iPSC lines

We used human iPSC lines derived from dermal fibroblasts from six healthy individuals with no known disease. All lines were obtained from the publicly available iPSCORE collection, developed by Dr. Kelly A. Frazer at the University of California, San Diego, as part of the NHLBI Next Generation Consortium (23). The lines are distributed by the WiCell Research Institute (Madison, WI, USA) or are available upon request from Dr. Frazer.

The six iPSC lines were derived from three females and three males of Asian ancestry, with age at donation ranging from 18 to 30 years. Detailed metadata for each individual is listed below: Individual 1 (UCSD129i-75-1/iPSCORE_75_1): Female, age 30, Asian (Irani); Individual 2 (UCSD132i-78-1/iPSCORE_78_1): Female, age 21, Asian (Chinese); Individual 3 (UCSD143i-87-1/iPSCORE_87_1): Female, age 21, Asian (Chinese); Individual 4 (UCSD178i-17-3/iPSCORE_17_3): Male, age 18, Asian (Japanese); Individual 5 (UCSD138i-84-1/iPSCORE_84_1): Male, age 21, Asian (Chinese); Individual 6 (UCSD154i-90-1/iPSCORE_90_1): Male, age 23, Asian (Chinese).

### iPSC maintenance and cardiomyocyte differentiation

iPSCs were cultured at 37 °C in a humidified atmosphere with 5% CO_2_ and ambient oxygen. Cells were maintained on Matrigel hESC-qualified Matrix (354277, Corning, Bedford, MA, USA) at a 1:100 dilution in mTeSR1 medium (85850, StemCell Technologies, Vancouver, BC, Canada), supplemented with 1% Penicillin-Streptomycin (30-002-Cl, Corning). Cultures were passaged every 4 - 6 days using a dissociation reagent containing 0.5 mM EDTA and 300 mM NaCl in PBS when they reached 70 - 80% confluence.

Directed differentiation of iPSCs into cardiomyocytes was performed as previously described (22). iPSCs were seeded in 10 cm Matrigel-coated dishes and cultured to 85 - 90% confluence. On Day 0, differentiation was initiated by replacing the culture medium with Cardiomyocyte Differentiation Medium (CDM) containing 12 μM CHIR99021 trihydrochloride (4953, Tocris Bioscience, Bristol, UK). The CDM formulation included 500 mL RPMI 1640 (15-040-CM, Corning), 10 mL B-27 minus insulin supplement (A1895601, ThermoFisher Scientific, Waltham, MA), 5 mL GlutaMAX (35050-061, ThermoFisher Scientific), and 5 mL Penicillin-Streptomycin. After 24 hours (Day 1), the media was replaced with fresh CDM. On Day 3, cells were treated with 2 μM Wnt-C59 (5148, Tocris Bioscience) in CDM to inhibit Wnt signaling. CDM was replaced on Days 5, 7, 10, and 12 to support continued cardiomyocyte differentiation.

On Day 14, cardiomyocytes were metabolically selected for using glucose-free, lactate-supplemented Purification Media. Purification Media consists of 500 mL RPMI without glucose (11879, ThermoFisher Scientific), 106.5 mg L-ascorbic acid 2-phosphate sesquimagnesium salt (sc228390, Santa Cruz Biotechnology, Santa Cruz, CA), 3.33 mL of 75 mg/mL human recombinant albumin (A0237, Sigma-Aldrich, St. Louis, MO), 2.5 mL of 1 M sodium lactate in HEPES buffer (L7022, Sigma-Aldrich), and 5 mL Penicillin-Streptomycin. Purification Media was exchanged on Days 16 and 18. On Day 20, iPSC-CMs were dissociated using 0.05% Trypsin-EDTA (25-053 Cl, ThermoFisher Scientific) for 10 - 15 min and neutralized with Cardiomyocyte Maintenance Medium (CMM), composed of 500 mL glucose-free DMEM (A14430-01, ThermoFisher Scientific), 50 mL FBS (S1200-500, Genemate), 990 mg galactose (G5388, Sigma-Aldrich), 5 mL 100 mM sodium pyruvate (11360-070, ThermoFisher Scientific), 2.5 mL 1 M HEPES (SH3023701, ThermoFisher Scientific), 5 mL GlutaMAX, and 5 mL Penicillin-Streptomycin. Cells were sequentially filtered through 100 μm and 40 μm nylon mesh strainers to generate a single-cell suspension.

iPSC-CMs were quantified and plated onto 0.1% gelatin-coated culture plates in CMM media. 50,000 iPSC-CMs were plated per well of a 96-well plate for viability assays; 600,000 cells per well of a 12-well plate for immunofluorescence staining, and 600,000 cells per well of a 12-well plate for RNA-seq. Cells were matured in culture for an additional 10 days, with media replaced on Days 23, 25, 27, 28, and 30.

### Quantification of iPSC-CM purity

Cardiomyocyte differentiation efficiency for each individual was assessed by measuring the expression of cardiac troponin T (TNNT2) by flow cytometry. Day 27 - 31 iPSC-CMs were dissociated with 0.05% Trypsin-EDTA to generate a single-cell suspension. One million cells per sample were stained with Zombie Violet Fixable Viability Dye (423113, BioLegend, San Diego, CA, USA) for 15 min at 4 °C in the dark, followed by fixation and permeabilization using the FOXP3/Transcription Factor Staining Buffer Set (00-5523, ThermoFisher Scientific) for 30 min at 4 °C. Fixed and permeabilized cells were stained with 5 µL of PE-conjugated anti-cardiac Troponin T antibody (Clone 13-11, 564767, BD Biosciences, San Jose, CA, USA) diluted in permeabilization buffer and incubated for 45 min at 4 °C in the dark. After three washes in permeabilization buffer, cells were resuspended in autoMACS Running Buffer (130-091-221, Miltenyi Biotec, Bergisch Gladbach, Germany) for acquisition. Control samples included iPSC-CMs that were unlabeled, iPSC-CMs labeled with TNNT2 only, and iPSC-CMs labeled with viability stain only. Flow cytometry was performed on a FACSymphony Cell Analyzer (BD Biosciences). 10,000 events were collected per sample. A set of live cells, based on the viability stain, was obtained following removal of debris and selection of singlets. iPSC-CM purity was quantified as the proportion of live, TNNT2-positive cells relative to the total live population.

### Drug preparation

CX-5461 (50-196-9852; MedChem Express, Monmouth, NJ, USA) was reconstituted in DMSO to generate a 1 mM stock solution and stored at -80 °C. Doxorubicin hydrochloride (D1515; Sigma-Aldrich) was dissolved in DMSO to a 10 mM stock and stored at -80 °C. Working concentrations for both compounds were prepared in CMM media immediately prior to treatment. DMSO (vehicle; VEH) controls were volume-matched to the highest corresponding drug concentration for each treatment.

### Cell viability assay

Viability assays were performed using iPSC-CMs from three individuals (Individuals 1, 2, 3) on Day 27 ± 1 of differentiation. iPSC-CMs from each individual were treated on one 96-well plate in quadruplicate with VEH (DMSO, 0 µM) and a range of concentrations (0.1 - 10 µM) for CX and DOX, as well as a 50 µM concentration for CX. Four wells contained untreated iPSC-CMs and four wells contained CMM media only. Fluorescent intensity was measured at 0 h and 48 h following treatment using the PrestoBlue Cell Viability Reagent (A13262; Invitrogen), according to the manufacturer’s instructions. Fluorescence was measured using a Synergy H1 plate reader (BioTek) at 560 nm excitation and 590 nm emission.

Fluorescence values were averaged across quadruplicate wells at each timepoint. The four media-only wells were averaged and denoted as background fluorescence at each timepoint. Timepoint-matched background fluorescence values were subtracted from VEH and drug-treated sample fluorescence values. These background-corrected fluorescence values were divided by the untreated background-corrected fluorescence values. Normalized, background-corrected fluorescence values at 48 h were divided by normalized, background-corrected fluorescence values at 0 h to obtain viability measurements for each treatment. Dose response curves were fitted for each drug in each individual using viability values and a four-parameter log-logistic regression model (LL.4) as implemented in the drc package (v3.0-1) in R (24). The half-maximal lethal dose (LD50) was extracted from the fitted models. LD50 values were compared across drugs using a paired two-tailed t-test. Viability effects were also determined at individual drug concentrations (0.1, 0.5, 1, 5, 10 µM) by comparing to the VEH (DMSO, 0 µM) samples using paired t-tests. *P* < 0.05 is considered significant.

### γ-H2AX immunofluorescence staining and quantification

Day 27-31 iPSC-CMs were treated with 0.1, 0.5, 5 or 10 µM CX, DOX, or VEH for three, 24, or 48 h. Following treatment, iPSC-CMs were fixed in 4% paraformaldehyde for 15 min at room temperature, then permeabilized with 0.25% DPBS-T (0.25% Triton X-100 in DPBS) for 10 min. After washing, cells were blocked with 5% BSA in DPBS-T for 30 min at room temperature and incubated overnight at 4 °C with anti-phospho-Histone H2A.X (Ser139) rabbit monoclonal antibody (1:500 dilution; NC1602516; Fisher Scientific) prepared in 1% BSA in DPBS-T. The following day, cells were washed and incubated for 1 h at room temperature with a donkey anti-rabbit Alexa Fluor 594-conjugated secondary antibody (1:1000 dilution; A-21207; Invitrogen). Nuclei were counterstained using Hoechst 33342 (PI62249; Thermo Scientific) for 10 min in the dark.

Immunofluorescence images were acquired under consistent exposure settings across conditions. One hundred nuclei were counted across two or three fields of view per treatment, and scored for presence or absence of γ-H2AX. Quantification of γ-H2AX-positive nuclei was performed using the Cell Counter plugin in ImageJ version 1.54i (25). The percentage of γ-H2AX– positive nuclei was calculated by dividing the number of positive nuclei by the total number of nuclei. The percentage of drug-treated iPSC-CMs positive for γ-H2AX was compared to the percentage in VEH-treated iPSC-CMs by a two-tailed paired t-test. *P* < 0.05 is considered significant.

### RNA extraction

Day 30 iPSC-CMs were treated with 0.1 or 0.5 µM CX, DOX or VEH for three, 24, or 48 h. Cells were flash-frozen and stored at -80 °C. Total RNA was extracted from flash-frozen iPSC-CMs using the RNeasy Mini Kit 250 (74106; QIAGEN, Germantown, MD, USA), following the manufacturer’s instructions. Extractions were performed in treatment- and timepoint-balanced batches of 12, with all conditions from a given individual processed in one batch. RNA concentration and integrity were assessed using the Agilent 2100 Bioanalyzer. All samples exhibited high-quality RNA with RNA Integrity Number (RIN) scores exceeding 8.0. Median RIN values across treatments for individuals 1-6 were 9.95, 9.75, 9.9, 9.05, 9.1, and 9.7, respectively.

### RNA-seq library preparation

Polyadenylated RNA was isolated from 150 ng of total RNA using the NEBNext Poly(A) mRNA Magnetic Isolation Module (E7490L, New England Biolabs, Ipswich, MA, USA), and RNA-seq libraries were prepared using the NEBNext Ultra II Directional RNA Library Prep Kit with Sample Purification Beads (E7765S; NEB) and indexed using NEBNext Multiplex Oligos for Illumina (96 Unique Dual Index Primer Pairs, E6440S). Libraries were prepared in two treatment- and timepoint-balanced batches consisting of all samples per individual for three individuals (Batch 1: Individuals 4-6; Batch 2: Individuals 1-3). Library quality was assessed using the Agilent 2100 Bioanalyzer. All 108 libraries were quantified, pooled, and sequenced across three AVITI paired-end sequencing runs (2 x 75 bp), yielding a median of 26,402,233 paired-end read pairs across samples.

### RNA-seq analysis

Raw sequencing reads were assessed for quality using MultiQC (v1.26) (26). Reads were aligned to the human reference genome (GRCh38/hg38) using the Rsubread package (v2.16.1) (27), and gene-level counts were quantified using the featureCounts function with built-in exon-based annotations (28). Count matrices were imported into R for downstream processing. All custom analysis scripts used for this project are available at https://github.com/mward-lab/Paul_CX_2025 made possible by the workflowr package (29).

Counts were transformed into log_2_ counts per million (log_2_cpm) using the cpm() function from the edgeR package (v4.0.16) (30). We excluded genes with a mean log_2_cpm ≤ 0 across all 108 samples, yielding 14,279 expressed genes for downstream analysis.

### Gene expression correlation between iPSC-CMs and human tissues

Gene expression data across 24 human tissues were obtained from Jiang *et al*. (31). We calculated the median log_2_cpm expression across all VEH-treated iPSC-CM samples. Pearson correlations were computed between VEH iPSC-CMs and each tissue using the 14,279 genes expressed in iPSC-CMs. Correlation values were visualized as a heatmap using the ComplexHeatmap package (v2.18.0) (32).

### Principal component analysis

We performed principal component analysis (PCA) on log_2_cpm values using the prcomp() function in the R Stats package (v4.3.0) (33), with log_2_cpm values centered prior to analysis. The top principal components (PCs) were visualized using ggfortify (v0.4.17) (34). Five covariates in the study: individual, sex, drug treatment, drug concentration and treatment time were assessed for association with the top PCs using a linear model. *P*-values were derived from the F-statistic, where *P* < 0.05 is considered significant.

### Differential gene expression analysis

We identified differentially expressed genes (DEGs) across all treatment conditions using the edgeR-voom-limma pipeline (35) and raw counts from the 14,279 expressed genes. Raw counts were normalized using the trimmed mean of M-values (TMM) method to account for differences in library size and composition and transformed with voom to estimate and account for the mean– variance relationship. DEGs were determined by linear modeling and empirical Bayes moderation. CX and DOX treatments were contrasted against the time- and concentration-matched VEH samples. Individual was treated as a random effect using the duplicate correlation function. Genes with an adjusted *P* < 0.05 (Benjamini–Hochberg correction) are classified as differentially expressed.

### Gene expression cluster analysis

We jointly modeled gene expression responses using the Cormotif package in R (v1.48.0) (36), which implements a Bayesian clustering approach to identify shared expression patterns, or correlation motifs, across multiple comparisons. TMM-normalized log_2_cpm values were used as input. We paired each drug treatment with its corresponding VEH control at three, 24, and 48 h. Separate models were constructed for the 0.1 µM and 0.5 µM datasets, each encompassing six pairwise comparisons. For both concentrations, we fit models across a range of motif numbers (K = 1 to 8) and selected the optimal model based on Bayesian information criterion (BIC) and Akaike information criterion (AIC). Following model fitting, we extracted gene-wise posterior probabilities of differential expression from the best-fit Cormotif model for each concentration. Genes were classified into discrete clusters based on their joint probability patterns. Genes with posterior probabilities < 0.5 across all treatments were designated as the Non-response cluster for both drug concentrations. For the 0.1 µM concentration: the CX_DOX_1 cluster is defined as genes with posterior probability > 0.5 in DOX_24hr, DOX_48hr, CX_24hr, CX_48hr, and p < 0.5 in DOX_3hr and CX_3hr. The DOX-specific_1 cluster is defined by posterior probability > 0.5 in DOX_24hr, DOX_48hr, and p < 0.5 in DOX_3hr and all CX timepoints. For the 0.5 µM concentration: the CX_DOX_2 cluster is defined by posterior probability > 0.5 in CX_24hr, CX_48hr, and p ≥ 0.1 for CX_3hr, p ≥ 0.02 for DOX_3hr, p < 0.5 for DOX_24hr and DOX_48hr. The CX_DOX_3 cluster has p > 0.5 in DOX_24hr, DOX_48hr, CX_24hr, CX_48hr, 0.02 ≤ p < 0.5 for DOX_3hr and p < 0.5 CX_3hr. DOX-specific_2 has p > 0.5 in DOX_3hr, DOX_24hr, p ≥ 0.02 in DOX_48hr and p < 0.5 in all CX timepoints. DOX-specific_3 has p > 0.5 for DOX_24hr, DOX_48hr, p < 0.5 for CX_3hr, CX_24hr, CX_48hr, DOX_3hr.

### Comparison of DEGs with CX DEGs in colorectal cancer cells

We obtained RNA-seq data from colorectal cancer cells treated with 0.5 µM CX or VEH for six, 24, or 48 h (4). First, we compared the log_2_ fold change values for the colorectal cancer cells with our data in iPSC-CMs. We calculated the Pearson correlation coefficient across all drug-VEH comparisons. Second, we obtained the set of DEGs (FDR < 0.05) in colorectal cancer cells at each time point. We combined the DEGs across timepoints to obtain a set of 1,228 unique DEGs. We calculated the proportion of colorectal cancer CX DEGs amongst our set of DEGs across treatments. Chi-square tests were used to compare overlap proportions between CX and DOX treatments at matched concentrations and timepoints. *P* < 0.05 is denoted as significant.

### Gene Ontology enrichment analysis

We performed functional enrichment analysis for both DEGs and response clusters. For each gene set, enrichment of GO terms in the Biological Process (BP) category was performed using the clusterProfiler package (v4.10.1) (37) in R. G-quadruplex terms were obtained from the Molecular Function category. The background gene set consisted of all expressed genes in the dataset (n = 14,279). GO terms with FDR-adjusted *P* < 0.05 are considered significant.

### DNA Damage Response gene expression analysis

We obtained 69 DNA damage-associated genes from the Molecular Signatures Database (MSigDB v7.5.1) (38). The effect sizes (log_2_ fold change) from the differential expression analysis across treatments were visualized for the 65 genes expressed in our data.

### p53 target gene expression analysis

We obtained a set of 346 p53 target genes (39). The effect sizes (log_2_ fold change) from the differential expression analysis across treatments were visualized for the 300 genes expressed in our data. Effect sizes across treatments were compared between concentration- and drug-matched CX and DOX treatments. *P* < 0.05 is considered significant.

### Heart-specific gene enrichment analysis

We obtained a set of 419 genes that are specifically expressed in the heart based on transcriptomic data (40). Genes were defined as heart-specific if they exhibited a ≥ 4-fold higher mRNA expression in heart tissue compared to all other human tissues. The proportion of heart-specific genes that overlap with response gene clusters was calculated. A Fisher’s exact test was used to compare enrichment between each response cluster and the Non-response cluster at each drug concentration. *P* < 0.05 is considered significant.

### Tissue-specificity analysis

We obtained heart left ventricle tissue-specificity scores from GTEx RNA-seq data (31). We collated tissue-specificity scores for all genes in each response cluster. We compared scores between the Non-response cluster and response clusters using Wilcoxon rank-sum tests, where significant differences are denoted when *P* < 0.05.

### Gene expression response in anthracycline-induced cardiotoxicity loci

We curated a set of DOX-induced cardiotoxicity genes from three previously published studies as described below. We obtained the 50 SNPs most significantly associated with anthracycline-induced cardiotoxicity from a GWAS study involving ∼3,000 breast cancer patients treated with anthracyclines (41) as well as 14 SNPs previously associated with anthracycline-induced cardiotoxicity and recommended for pharmacogenomic testing in pediatric patients by the Canadian Pharmacogenomics Network for Drug Safety (42). SNPs were mapped to the nearest transcription start site using BEDtools to obtain a gene set (43). We also collated all genes whose expression is associated with these SNPs in any human tissue using the GTEx v10 eQTL database (44). We removed genes not expressed in our dataset and filtered out redundant entries. We also incorporated genes prioritized by transcriptome-wide association analysis (TWAS) for their role in anthracycline-induced cardiotoxicity (45), and anthracycline-induced cardiotoxicity genes that have been functionally-validated in iPSC-CMs (46). The effect sizes (log_2_ fold change) from the differential expression analysis across treatments were visualized for this gene set.

### Gene expression response in heart failure-associated loci

We obtained a set of 273 loci associated with heart failure through the GWAS catalog (47), and extracted the mapped genes. 118 unique mapped genes are expressed in our data. The effect sizes (log_2_ fold change) from the differential expression analysis across treatments were visualized for the set of top 30-associated genes. Effect sizes across treatments were compared between concentration- and timepoint-matched CX and DOX treatments. *P* < 0.05 is considered significant.

## Results

### CX does not induce DNA damage or loss of viability in iPSC-CMs at sub-micromolar concentrations

To assess the cardiotoxic potential of CX relative to DOX, we used an *in vitro* model based on iPSCs derived from six healthy individuals (three females and three males) with no known cardiac disease or cancer diagnosis (Figure 1A). We differentiated the six iPSC lines into beating cardiomyocytes. The majority of iPSC-CMs from each individual express the cardiac-specific marker, cardiac troponin T, as measured by flow cytometry (median purity = 98.4%; Figure S1).

**Figure 1:**
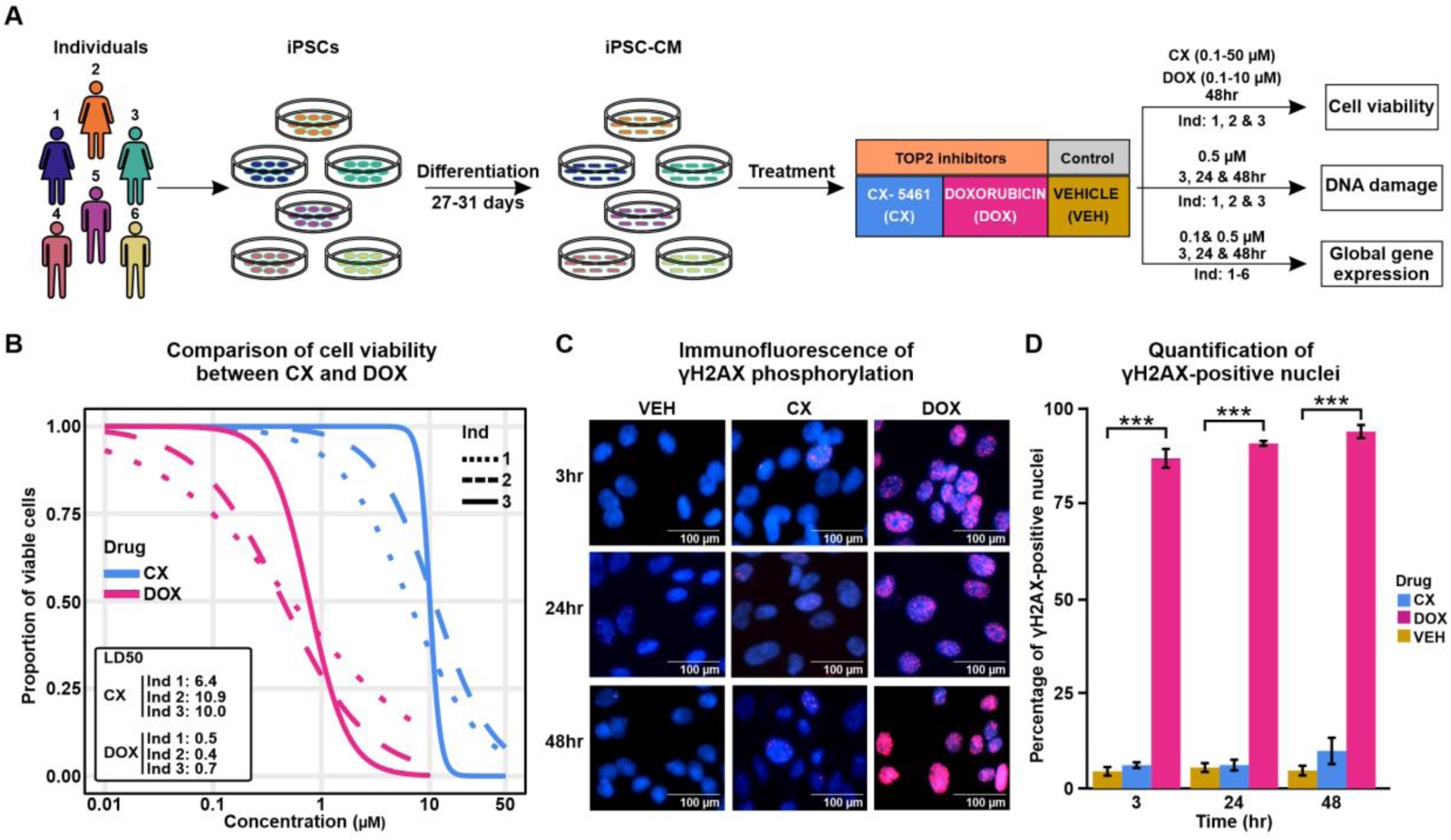
CX induces minimal cytotoxicity and DNA damage in iPSC-CMs compared to DOX. **(A)** Experimental design of the study. iPSCs derived from three males and three females were differentiated into cardiomyocytes (iPSC-CMs) and treated with TOP2 inhibitor drugs CX-5461 (CX) or Doxorubicin (DOX) or a vehicle control (VEH) for three, 24 or 48 hours. Effects on cell viability, DNA damage, and gene expression were measured. **(B)** Proportion of viable cardiomyocytes following exposure to increasing concentrations of CX (blue) or DOX (pink) for 48 hours in three individuals. Viability data were collected in quadruplicate for each drug concentration, normalized and expressed relative to pre-treatment values. Dose response curves derive from mean viability measurements fit with a four-parameter logistic regression. **(C)** Immunostaining of the DNA damage marker, γ-H2AX (red), and Hoechst nuclear stain (blue) in iPSC-CMs from a representative individual (Individual 1) treated with 0.5 µM CX, DOX or VEH over time. Scale bar: 100 μm. **(D)** Percentage of VEH-, CX- and DOX-treated iPSC-CMs that stain positive for γ-H2AX. Data representative of 100 cells per treatment per individual. Data are presented as mean ± SD across three individuals (1,2,3). Asterisk represents a statistically significant difference in γ-H2AX expression (\*\*\**P* < 0.001).

To evaluate drug-induced cytotoxicity, we treated iPSC-CMs from three individuals with a range of concentrations of CX (0 - 50 µM) and DOX (0 - 10 µM) and a vehicle (VEH) control for 48 hours and assessed cell viability using a resazurin-based fluorometric assay. Both drugs induced dose-dependent decreases in cell viability across individuals. However, DOX is approximately 20 times more cytotoxic than CX (median LD50 DOX= 0.53 µM, median LD50 CX= 10.03 µM; two-tailed paired t-test; *P* = 0.02; Figure 1B). The concentration of CX in plasma from patients treated with the drug range from 0.5 – 3.5 µM (6). We therefore selected two sub-micromolar, clinically-relevant concentrations of CX, 0.1 and 0.5 µM, that have been used in prior *in vitro* studies for further characterization (5, 14). There is no significant difference in viability between VEH and treatment with 0.1 or 0.5 µM CX or DOX (Figure S2). To investigate drug effects over time, we treated cells with either 0.1 or 0.5 µM CX and DOX or VEH for three, 24 or 48 hours.

Both DOX and CX have been reported to induce cytotoxicity by initiating DNA damage in cancer cells. While DOX has also been shown to induce DNA damage in iPSC-CMs, the effects of CX have not been tested. To investigate whether CX has genotoxic effects at sub-lethal concentrations, we quantified the expression of γ-H2AX, a marker of DNA double-strand breaks, in drug-treated iPSC-CMs by immunofluorescence staining. Following treatment with 0.1 µM DOX there is an increase in γ-H2AX staining compared to VEH with 91.8% of cells staining positive for γ-H2AX as early as three hours following treatment (t-test; *P* < 0.05 for all timepoints; Figure S3 A-B). In contrast, CX treatment results in similar levels of γ-H2AX as VEH across timepoints with at most 6.1% of cells staining positive for the double-strand break marker. Treatment with a higher concentration of CX (0.5 µM) does not increase γ-H2AX staining compared to VEH at any time point (Figure 1 C-D). The proportion of γ-H2AX-positive cells is low even following 24 hours of CX treatment at ten to 100 times the concentration (5 µM = 5.3%, 10 µM = 9.4%; Figure S4). As expected, there is an increase in γ-H2AX in cells treated with 0.5 µM DOX compared to VEH at all timepoints (*P* < 0.05; Figure 1C-D). These results demonstrate that treatment with CX results in minimal DNA damage and loss of viability in cardiomyocytes compared to DOX.

### CX elicits a distinct and attenuated transcriptomic response compared to DOX

To characterize the impact of CX treatment on the transcriptome of human cardiomyocytes, we performed bulk RNA sequencing on iPSC-CMs from six individuals treated with 0.1 and 0.5 µM CX, DOX, or VEH for three, 24, and 48 hours, yielding data from 108 samples (Table S1). Sequencing metrics indicated high data quality across all treatments, with a median yield of 26,402,233 read pairs across samples, and uniform read depth across treatment, concentration, and timepoint groups (Figure S5 A-C; Table S2). Following genome alignment, counting of reads mapping to genes, and removal of lowly expressed genes, we obtained a final set of 14,279 expressed genes.

To determine the physiological relevance of our *in vitro* human cardiomyocyte model to human heart tissue, we correlated our iPSC-CM gene expression data (median VEH log_2_cpm) with gene expression data from human tissues (31). Amongst the 24 tissues tested, iPSC-CMs are most similar to heart tissue (r = 0.81 for right atrium, r = 0.79 for left ventricle; Figure S6A). Similarly, iPSC-CMs express a range of cardiac genes including *TTN*, *MYH6*, *ACTN2*, and *RYR2* (Figure S6B). When considering the gene expression data from all 108 samples, the data clusters primarily based on whether the iPSC-CMs were treated with DOX for 24 or 48 hours or not (Figure S7). Principal component analysis reveals that PC1, accounting for 31.6% of the variance, associates with drug type, drug concentration and treatment time (Figure 2A and Figure S8). The second principal component, representing 15.6% of the variation is associated with sex and individual indicating that the primary sources of variation in the data associate with known biological factors (Figure S8).

**Figure 2:**
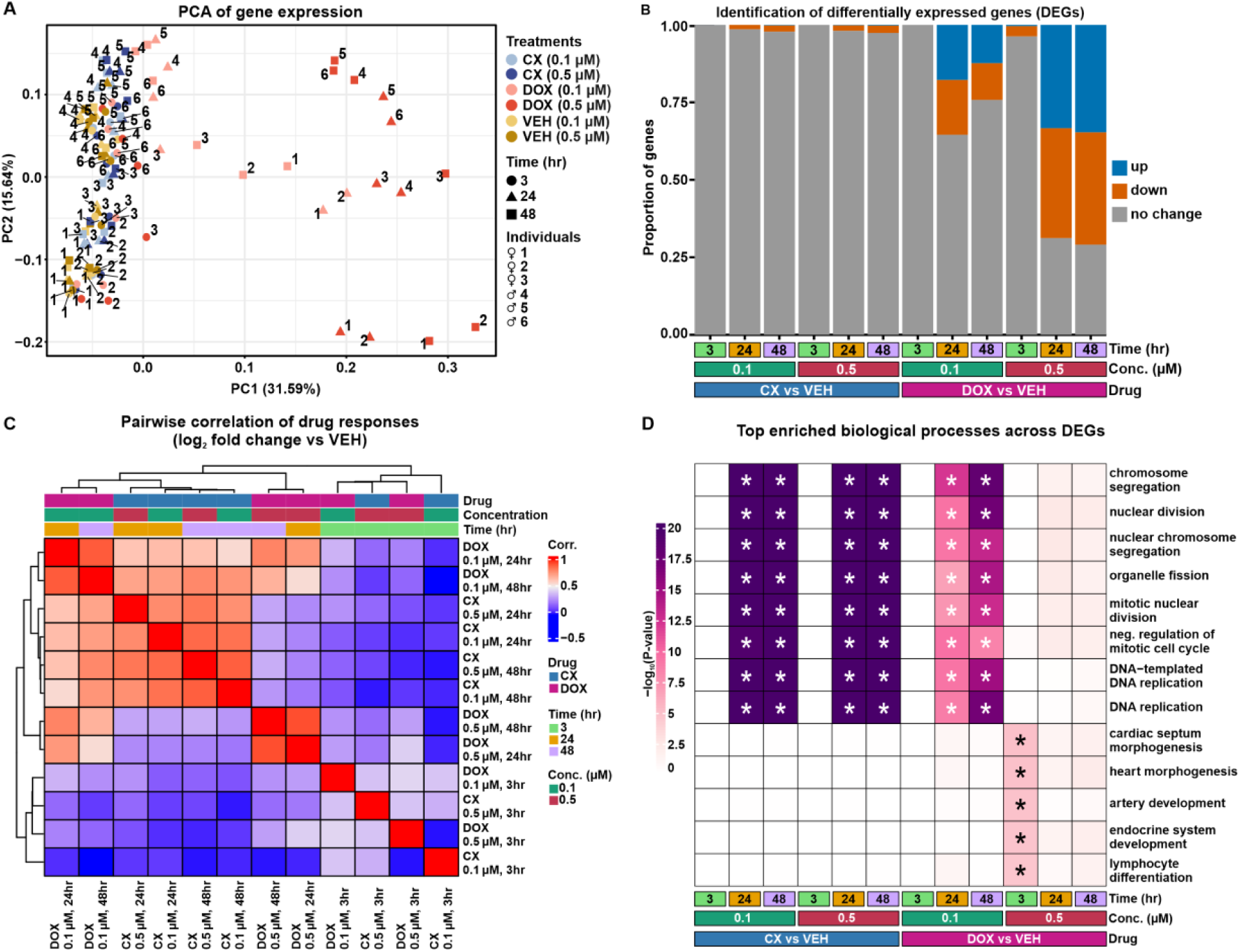
CX induces minimal effects on the iPSC-CM transcriptome compared to DOX. **(A)** Gene expression profiles (log_2_cpm) across 14,279 expressed genes from 108 samples representing six individuals (1,2,3,4,5,6), two sexes (male, female), three treatments (DOX (red), CX (blue), VEH (yellow)), two drug concentrations (0.1 µM (pale shade), 0.5 µM (dark shade)), and three treatment times (three hours: circles; 24 hours: triangles; 48 hours: squares). **(B)** Proportion of differentially expressed genes (DEGs) between drug (CX and DOX) and VEH at either 0.1 or 0.5 µM drug concentrations following three, 24 or 48 hours of treatment amongst all expressed genes. DEGs are categorized by whether they are upregulated in response to drug (up) or downregulated in response to drug (down). **(C)** Correlation between drug responses across drugs. The log_2_ fold change of expression between each drug treatment and VEH was calculated for all 14,279 expressed genes at each concentration and timepoint. Values were compared across all treatment groups by hierarchical clustering of the pairwise Pearson correlation values. Drugs are colored by type (CX: blue; DOX; pink), concentration (0.1 µM: teal; 0.5 µM: magenta), and treatment time (three hours: light green; 24 hours: orange; 48 hours: purple). **(D)** Top biological processes enriched across DEGs (adj. *P* < 0.05). The top five biological processes enriched in each drug treatment were collated, and the unique set interrogated across DEGs. Enrichment in each treatment is represented as -log_10_ *P* values. Asterisk represents processes significantly enriched in a set of DEGs (adj. *P* < 0.05).

To quantify the impact of drug treatment on the iPSC-CM transcriptome, we identified differentially expressed genes (DEGs) between each drug treatment and the time-matched VEH (adj. *P* < 0.05; Figure 2B; Figure S9; Table S3-14). CX treatment induced hundreds of DEGs across drug concentrations and treatment time (**3 hour:** 0.1 µM CX vs VEH n = 1; 0.5 µM CX vs VEH n = 2; **24 hour:** 0.1 µM CX vs VEH n = 205; 0.5 µM CX vs VEH n = 278; **48 hour:** 0.1 µM CX vs VEH n = 318; 0.5 µM CX vs VEH n = 370). DOX treatment induced thousands of DEGs (**3 hour:** 0.1 µM DOX vs VEH n = 2; 0.5 µM DOX vs VEH n = 533; **24 hour:** 0.1 µM DOX vs VEH n = 5,064; 0.5 µM DOX vs VEH n = 9,828; **48 hour:** 0.1 µM DOX vs VEH n = 3,461; 0.5 µM DOX vs VEH n = 10,128). While DOX treatment results in a roughly even distribution of DEGs that are up- and downregulated at 24 and 48 hours, the majority of DEGs induced by CX at these timepoints are downregulated (93% of all CX DEGs and 51% of all DOX DEGs; Figure 2B and Figure S9).

To directly compare drug responses across drug types and time we calculated all pairwise correlations of effect sizes (log_2_ fold change). Samples separate primarily by whether they were treated for three hours or not, regardless of drug type or concentration (Figure 2C). Three-hour responses are generally weakly correlated consistent with the low number of DEGs at this timepoint (rho = 0.36 - 0.40). iPSC-CMs treated with 0.1 or 0.5 µM DOX or 0.5 µM CX are most similar in this group. Amongst the 24- and 48-hour treatment time samples, 0.5 µM DOX-treated samples cluster separately from the other treatments and are the most strongly correlated responses (rho = 0.91), followed by 0.1 µM DOX-treated samples (rho = 0.89). CX responses are closest to low dose DOX responses and show strong correlations amongst each other (rho = 0.75 – 0.82).

To gain insight into whether the gene expression changes induced by CX in iPSC-CMs are similar to those in cancer cells, we obtained DEGs and treatment effect sizes from colorectal cancer cells treated with 0.5 µM CX for six, 24 or 72 hours (4). Comparing our CX responses to responses in cancer cells revealed correlated effect sizes at the 24- and 48-hour timepoints (median pearson correlation = 0.22 - 0.34; *P* < 0.05; Figure S10A). Our DOX treatment effect sizes are also correlated with CX treatment in cancer cells albeit lower than that for CX (correlation = 0.13 - 0.22; *P* < 0.05; Figure S10A). These results suggest a degree of sharing of CX responses across cancer cells and cardiomyocytes, and across CX and DOX treatment. Indeed, we find that 33 - 40% of our 24- and 48-hour iPSC-CM CX DEGs are DEGs in at least one timepoint following CX treatment in colorectal cancer cells (n = 1,228 DEGs), and that the proportion is significantly higher than those overlapping DOX DEGs at these timepoints (Chi-square test; *P* < 0.05; Figure S10B).

We next asked which biological processes are enriched amongst DEGs compared to all expressed genes (Fishers’ exact test; adj. *P* < 0.05). The top processes enriched amongst CX DEGs relate to chromosome segregation and DNA replication (Figure 2D). These processes are enriched across drug concentrations at 24- and 48-hour timepoints, and are also amongst the most enriched in the 0.1 µM DOX-treated samples over time. Many biological processes are shared across CX and 0.1 µM DOX DEGs (Figure S11). Samples treated with 0.5 µM DOX for three hours are enriched for processes related to heart development, while there are no enriched processes at the later timepoints likely due to the large proportion of DEGs.

### CX induces a subset of DNA damage response genes but not RNA polymerase I-associated genes

We identified hundreds of biological processes enriched amongst CX DEGs. We next asked whether three broad functional categories previously associated with CX treatment in cancer cells are contained in this set. First, we considered that CX selectively inhibits RNA polymerase I-mediated transcription in cancer cells. We did not find any of the eight ‘transcription by RNA polymerase I’-associated terms enriched amongst CX or DOX DEGs suggesting cell-type specificity in this effect (Figure S12A). Second, we considered the ability of CX to stabilize G-quadruplex DNA structures and asked whether six G-quadruplex-associated terms are enriched in our data. The ‘G-quadruplex binding’ function is enriched in the set of DEGs identified post treatment with 0.5 µM CX at the 24- and 48- hour timepoints, suggesting that this activity is involved in iPSC-CMs as well as in previously identified cell types (Figure S12B). This function is not associated with any DOX DEGs. Third, we considered eight ‘DNA damage response’-associated terms. We found DNA repair and DNA damage signal transduction processes enriched amongst CX DEGs at 24- and 48-hour timepoints across concentrations (Figure S12C). These terms are also enriched amongst 0.1 µM DOX DEGs.

Given the enrichment of DNA damage response terms across CX DEGs, we determined the transcriptional response of a curated set of DNA damage response genes from the Molecular Signatures Database (38). As expected, DOX treatment affects the expression of many DNA damage response genes (Figure 3A). Upregulated genes include apoptotic genes (*FAS*, *CASP3* and *BBC3*), while downregulated genes include cell cycle checkpoint-related genes (*CCNB1, CDC25C* and *CDK1*). CX treatment also downregulates cell cycle genes, as well as DNA repair genes (*BRCA1*, *RAD51* and *CHEK2* for example) similar to DOX, but does not induce apoptotic gene expression.

**Figure 3:**
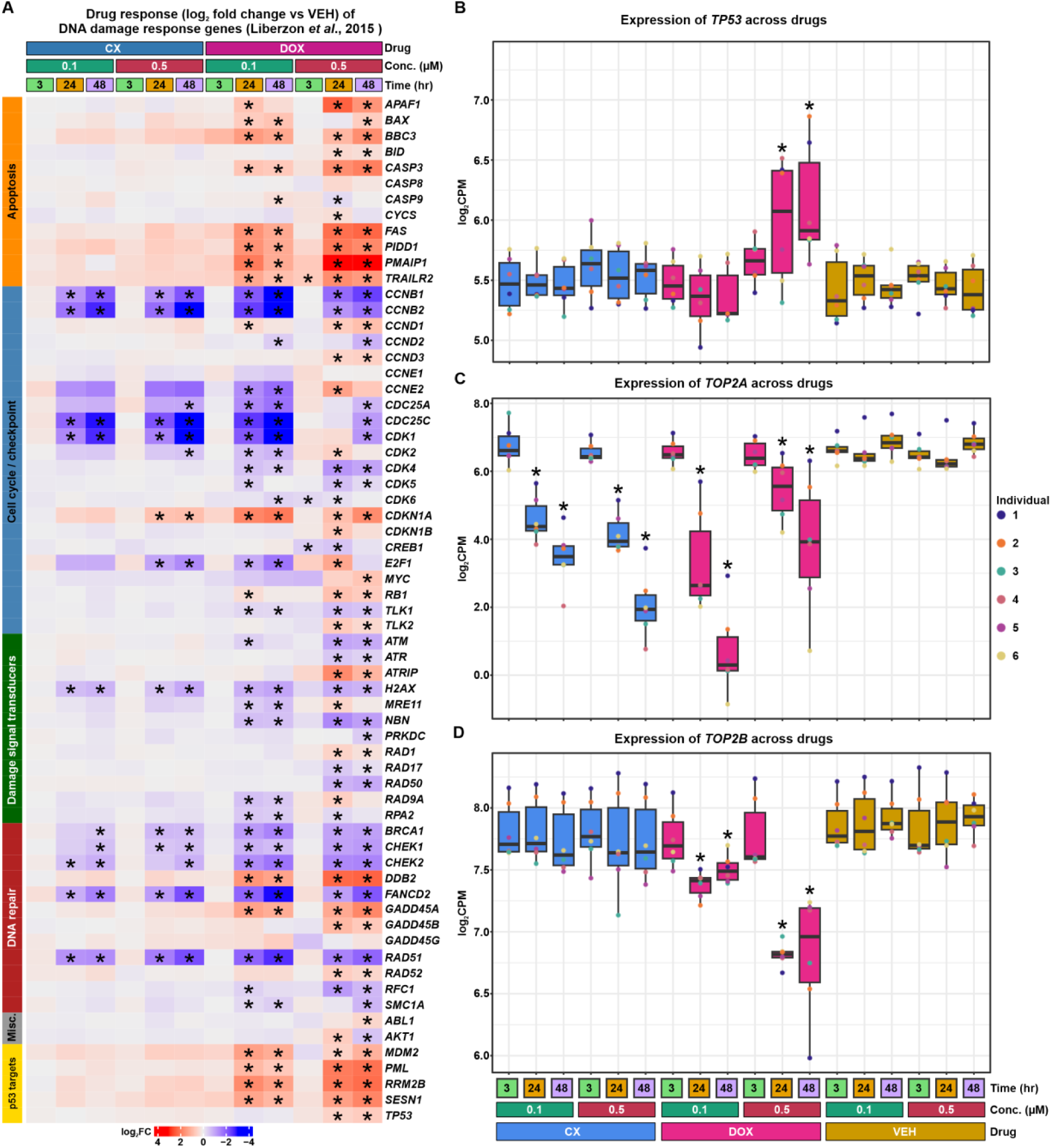
CX treatment affects DNA damage response gene expression. **(A)** Drug responses (log_2_ fold change) of DNA damage response genes (38). Asterisk represents genes that are classified as DEGs (adj. *P* < 0.05). **(B)** Expression of *TP53* (log_2_cpm) across CX, DOX and VEH treatments at two drug concentrations over time. Each treatment is represented by expression measurements from six individuals. Asterisk represents treatments where *TP53* is classified as a DEG (adj. *P* < 0.05). **(C)** Expression of *TOP2A* (log_2_cpm) as for (B). **(D)** Expression of *TOP2B* (log_2_cpm) as for (B).

p53 is a classic regulator of the DNA damage response. Its expression is induced following 24 and 48 hours of treatment with 0.5 µM DOX (Figure 3B). Neither CX, nor low concentrations of DOX, affect p53 gene expression suggesting a basis of the DNA damage response gene differences between drugs. Indeed, 24 and 48 hours of DOX treatment, at both concentrations, induces a stronger response on a curated set of 300 p53 target genes (39) than matched CX treatments (Wilcoxon rank-sum test; *P* < 0.05 Figure S13).

The DNA damage response induced by DOX in cardiomyocytes is TOP2B-dependent. TOP2B has recently been identified as the target of CX (14). We therefore asked whether expression of *TOP2B,* and its related isoform *TOP2A,* are affected by CX and DOX treatment in iPSC-CMs. We find that DOX decreases *TOP2B* expression following 24 and 48 hours of treatment at both concentrations (Figure 3C). CX has no effect on *TOP2B* mRNA levels at any timepoint or concentration. However, both drugs decrease *TOP2A* expression following 24 and 48 hours of treatment at both drug concentrations (Figure 3D). This is in line with both drugs decreasing the expression of cell cycle regulators.

### CX treatment induces gene expression changes shared with DOX

To directly compare CX and DOX-induced transcriptional changes, we jointly modeled the data to assess gene expression trajectories over time in a concentration-dependent manner (See Methods). We determined that three clusters represent the predominant patterns in the data from the samples treated with 0.1 µM CX, DOX and VEH (Figure S14A), and five clusters represent the samples treated with 0.5 µM CX, DOX and VEH (Figure S14B). At 0.1 µM, the majority of genes have a low probability of being differentially expressed across timepoints for both CX and DOX, and are therefore classified as Non-response genes (n= 12,308; Figure 4A; Table S15). There are 1,551 genes that have a high probability of differential expression following 24 and 48 hours of DOX treatment only, which are denoted as the DOX-specific_1 cluster. There are no gene clusters that show CX-specific responses. Instead, there are 415 genes that respond following 24 and 48 hours of CX and DOX treatment, denoted as the CX-DOX_1 cluster.

**Figure 4:**
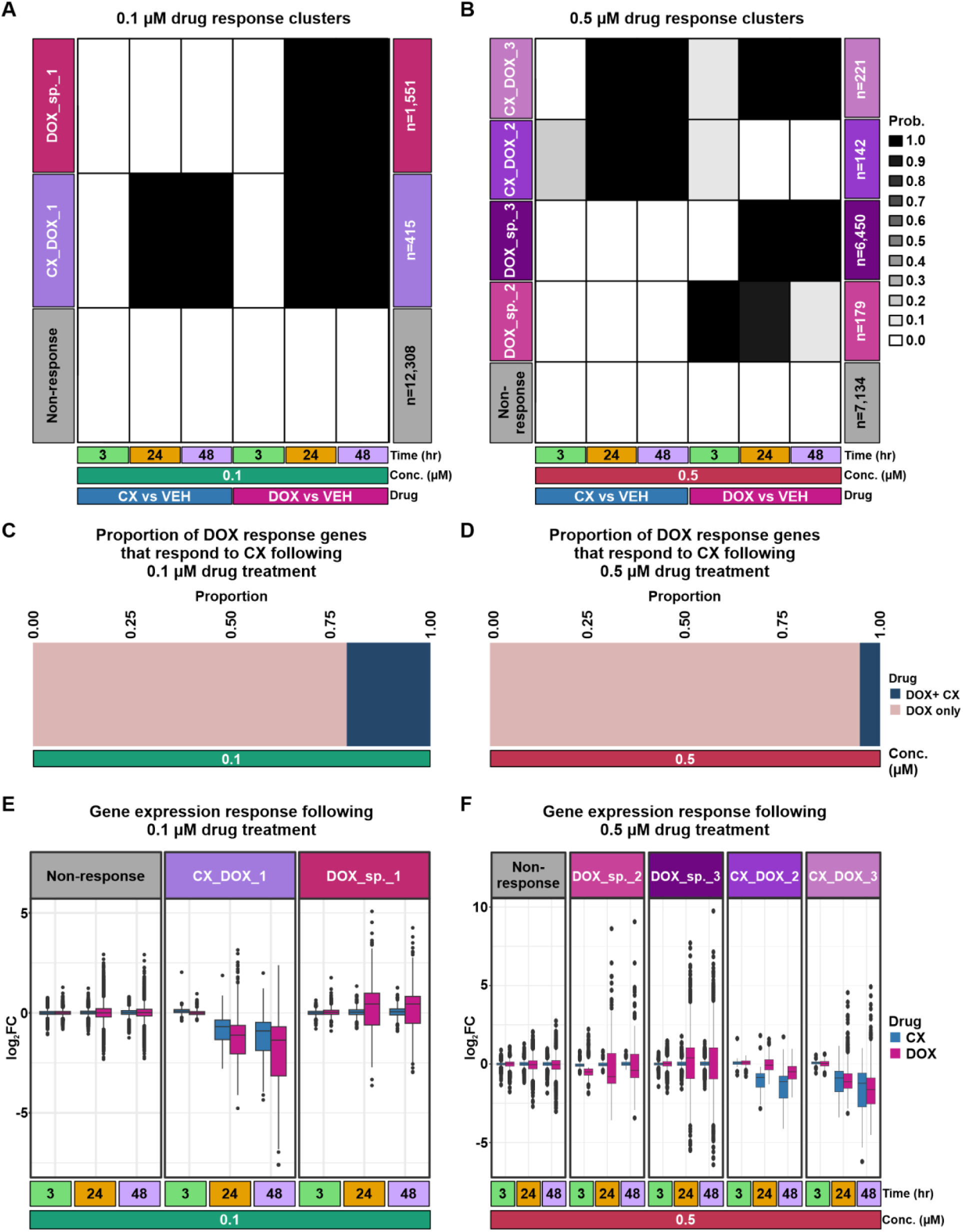
CX treatment induces gene expression changes shared with DOX. **(A)** Gene expression clusters following joint modeling of pairs of tests from the 0.1 µM drug-treated samples. Posterior probabilities of genes being differentially expressed following a treatment are represented by white to black shading. Genes are categorized by their posterior probability in each treatment. The grey cluster represents Non-response genes that have a low probability of differential expression across drugs. The lilac cluster represents genes that respond to CX and DOX. The magenta cluster represents genes that respond only to DOX. **(B)** Gene expression clusters following joint modeling of pairs of tests from the 0.5 µM drug-treated samples. Clusters are defined as described in (A). **(C)** The proportion of DOX response genes that also respond to CX following 0.1 µM drug treatment. **(D)** The proportion of DOX response genes that also respond to CX following 0.5 µM drug treatment. **(E)** The log_2_ fold change across CX and DOX treatments compared to VEH for genes in each cluster identified in the 0.1 µM drug-treated samples. **(F)** The log_2_ fold change across CX and DOX treatments compared to VEH for genes in each cluster identified in the 0.5 µM drug-treated samples.

Amongst the samples treated with 0.5 µM CX, DOX and VEH, there are 7,134 Non-response genes (Figure 4B; Table S16). Similar to the gene expression profiles from the 0.1 µM data, there are 6,450 genes that respond to DOX only at 24 and 48 hours (DOX-specific_3 cluster). The higher concentration of DOX results in an additional gene cluster including genes that respond strongly at three and 24 hours and weakly at 48 hours (DOX-specific_2; n = 179). At this higher concentration of drug there is still no CX-specific response. There is a cluster of genes that responds to CX at 24 and 48 hours and weakly to DOX at three hours and strongly at 24 and 48 hours (CX-DOX_3; n = 221), and another that responds weakly to CX at three hours and strongly at 24 and 48 hours and weakly to DOX at three hours (CX-DOX_2; n = 142). The proportion of DOX response genes that also respond to CX is greater in the 0.1 µM drug group than the 0.5 µM drug group (Chi-square test; *P* < 0.05; Figure 4C-D).

The genes in each cluster show the expected response to each drug based on the log_2_ fold change distributions between pairwise drug and VEH comparisons (Figure 4E-F). The Non-response groups show small drug effect sizes across drug concentrations, while the DOX-specific response clusters show increased absolute effect sizes in the DOX-treated groups. Clusters of genes with expression effects induced following both CX and DOX treatment tend to be downregulated in line with the majority of CX DE genes being downregulated. Individual genes in each of the response categories show the expected gene expression patterns (Figure S15). For example, *TRIP13* is a CX-DOX_1 response gene that is downregulated in response to both CX and DOX at 24 and 48 hours following 0.1 µM treatment (Figure S15A), while *BRCA1* is a CX-DOX_3 response gene that is downregulated at 24 and 48 hours following 0.5 µM treatment (Figure S15B). These results show that CX-induced effects on gene expression are shared with DOX at each timepoint. We next asked whether the genes that respond to 0.1 µM CX treatment also respond to 0.5 µM CX treatment. Of the 457 genes that respond to CX at either concentration, 321 (70%), are shared across concentrations suggesting robust effects on gene expression (Figure S16).

### CX response genes are not heart-specific

We next asked which biological processes are enriched amongst the eight drug response clusters. Similar to the pairwise DEG analysis, we find terms related to chromosome segregation and DNA repair amongst the response clusters shared between DOX and CX at both drug concentrations (adj. *P* < 0.05; Figure 5A; Figure S17). In contrast, the 0.5 µM DOX-specific_3 response cluster is enriched for terms including GPCR signaling and cardiac development.

**Figure 5:**
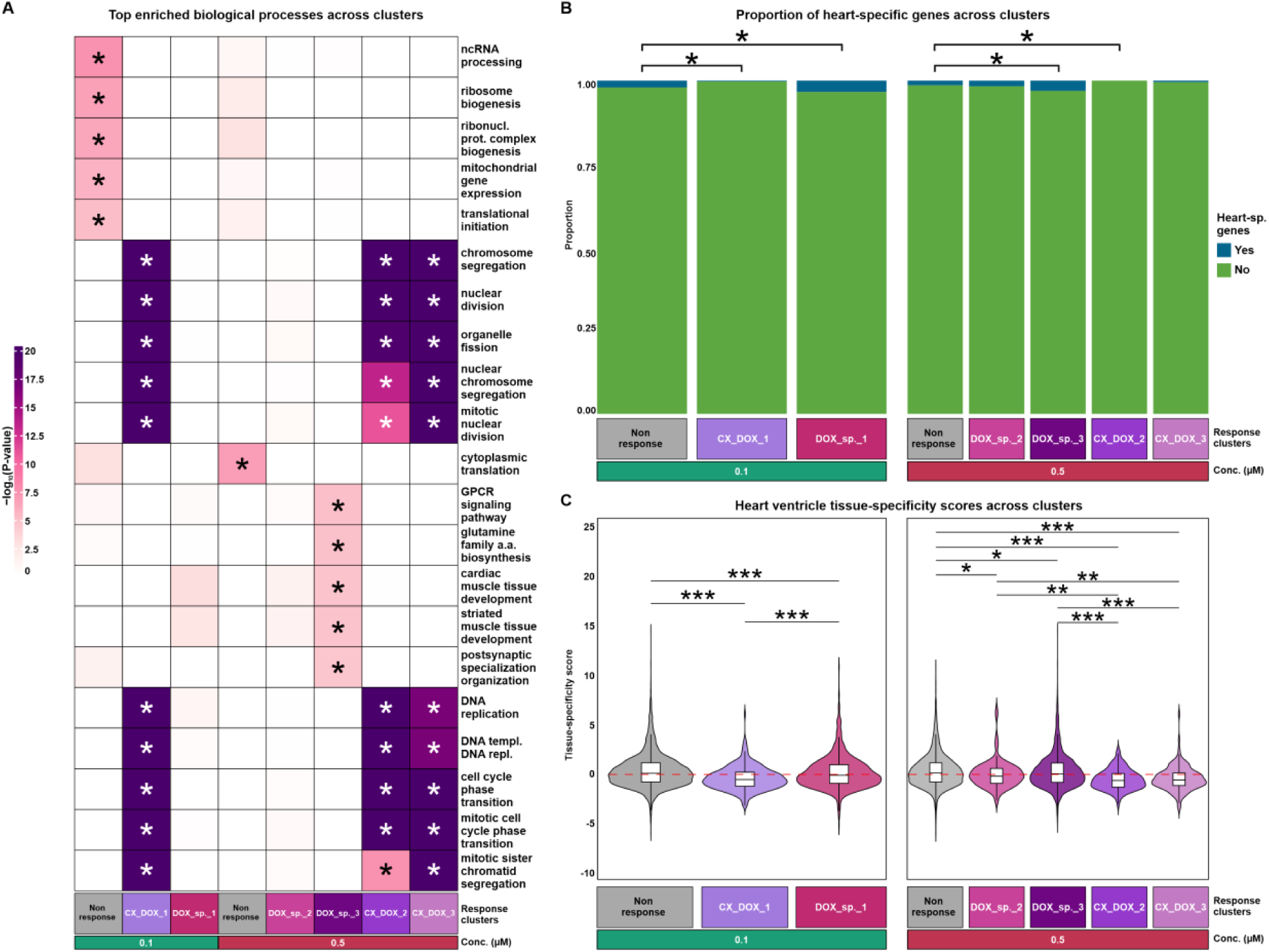
CX treatment induces minimal effects on heart-specific genes. **(A)** Top biological processes enriched across gene clusters (adj. *P* < 0.05). The top five biological processes enriched in each cluster were collated, and the unique set interrogated across clusters. Enrichment in each treatment is represented as -log_10_ *P* values. Asterisk represents processes significantly enriched in a cluster (adj. *P* < 0.05). **(B)** Proportion of heart-specific genes (48) amongst gene expression clusters. Asterisk represents a significant enrichment or depletion of heart-specific genes within a drug-associated gene expression cluster compared to the Non-response cluster. **(C)** Tissue-specificity scores of genes in each gene expression cluster. Tissue-specificity scores were obtained for each gene in left ventricle heart tissue (31). Higher tissue-specificity scores indicate genes that have higher specificity for heart tissue. Significant differences between clusters are shown (\**P* < 0.05, \*\**P* < 0.01, \*\*\**P* < 0.001).

Given the tissue-specific terms associated with DOX-specific clusters, we asked whether specific response clusters at each drug concentration are more likely to contain a set of 419 curated heart-specific genes (48) than the corresponding Non-response cluster. At 0.1 µM drug treatments, heart-specific genes are depleted amongst CX-DOX_1 cluster genes and enriched amongst DOX-specific_1 cluster genes (Fisher’s exact test; *P* < 0.05; Figure 5B). Similarly, at 0.5 µM drug treatments, the CX-DOX_2 cluster is depleted for heart-specific genes, and the DOX_specific_3 cluster is enriched.

We next considered tissue-specificity more broadly by comparing tissue-specificity scores for all genes expressed in the heart left ventricle across response clusters (31). At 0.1 µM drug treatments, tissue-specificity scores are significantly lower in both response clusters relative to the Non-response cluster (Wilcoxon rank-sum test; *P* < 0.05), indicating that these clusters contain genes that are broadly expressed across tissues (Figure 5C). However, the CX-DOX_1 cluster has a lower tissue-specificity score than the DOX-specific_1 cluster. Similarly, at 0.5 µM drug treatments, the CX-DOX_2 and CX-DOX_3 clusters have lower tissue-specificity scores than the Non-response cluster, and the DOX-specific_2 and DOX-specific_3 clusters (*P* < 0.05). Together, these results indicate that CX treatment does not preferentially affect the expression of heart-specific genes.

### CX does not affect the expression of genes in heart disease risk loci

DOX, the canonical anthracycline, is an effective anti-cancer drug. However, it can cause cardiotoxicity in some patients (49). DOX-induced cardiotoxicity is mediated through TOP2B (18). Given that both DOX and CX inhibit TOP2, we evaluated whether CX and DOX influence the expression of genes implicated in DOX-induced cardiotoxicity. We collated 72 anthracycline-induced cardiotoxicity risk genes from two GWAS (41, 42) and one TWAS (45) and assessed response to CX and DOX treatment. In line with prior studies, treating iPSC-CMs with DOX affects the expression of DOX-induced cardiotoxicity genes (22) (Figure S18). Of the 72 associated genes, 60 show expression changes following DOX treatment in at least one concentration and timepoint (adj. *P* < 0.05). In contrast, only one gene is affected by CX treatment. *GPSM2* is downregulated following treatment with both CX and DOX across concentrations at both 24 and 48 hours. We next focused on 25 DOX-induced cardiotoxicity genes that have been functionally validated (46). Eighty percent of these genes are classified as DOX DEGs (Figure 6A). None of these genes are CX DEGs at any timepoint or drug concentration.

**Figure 6:**
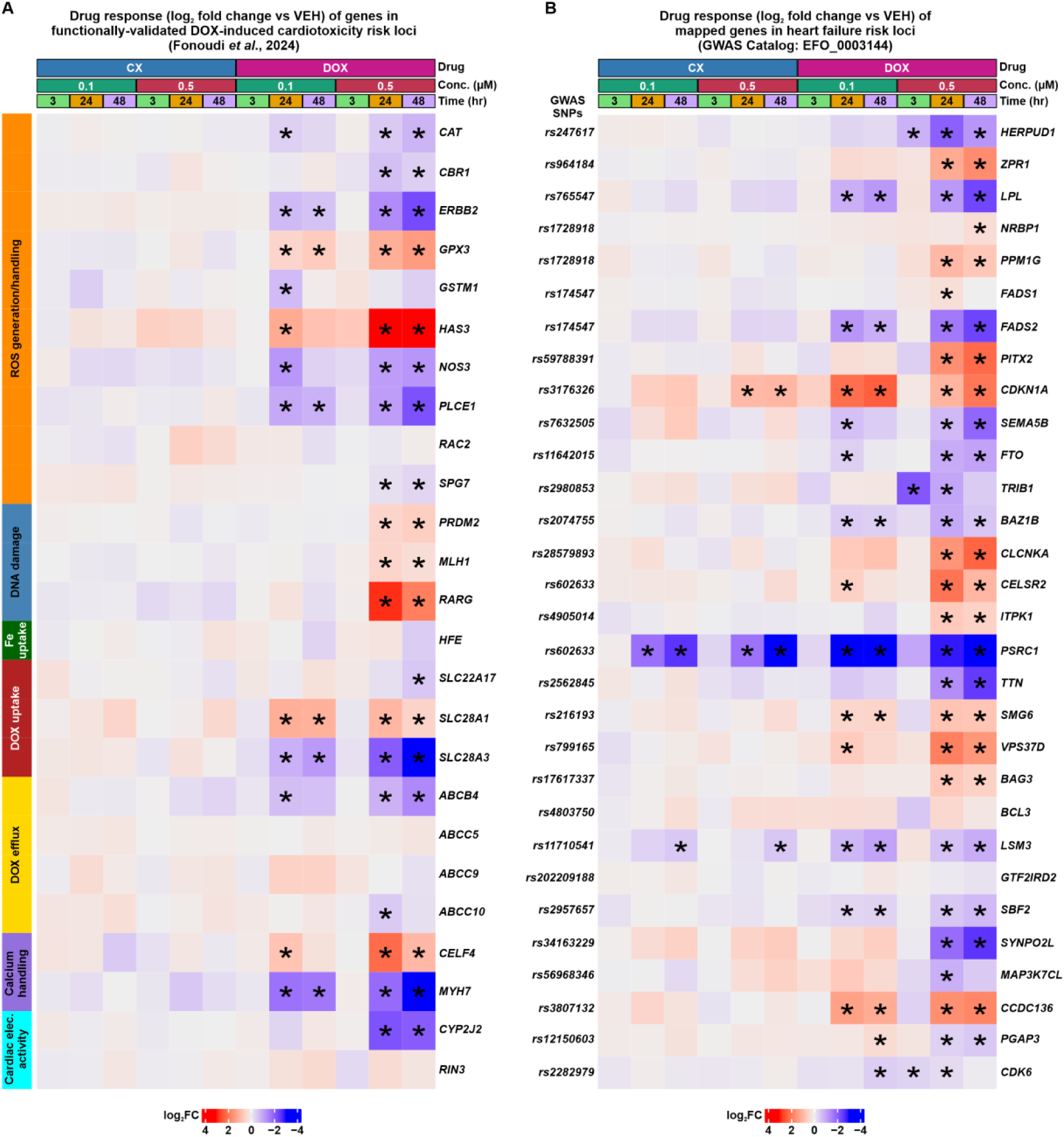
CX treatment does not affect the expression of genes in loci associated with DOX-induced cardiotoxicity. **(A)** Drug responses (log_2_ fold change) of functionally validated genes in DOX-induced cardiotoxicity loci (46). Asterisk represents genes that are classified as DEGs (adj. *P* < 0.05). **(B)** Drug responses (log_2_ fold change) of mapped genes in heart failure-associated loci (GWAS) (47). Mapped genes were obtained from the 30 SNPs most significantly associated with heart failure. Asterisk represents genes that are classified as DEGs (adj. *P* < 0.05).

Anthracyclines such as DOX, are increasingly recognized as having the potential to lead to heart failure (50, 51). We therefore examined the transcriptional responses of 118 genes associated with heart failure risk through GWAS across CX and DOX drug treatments (47). When considering the effect of the treatment (log_2_ fold change) at all 118 mapped risk genes at the highest concentration and timepoint, DOX induced a greater response amongst these genes than CX (Wilcoxon rank-sum test; *P* < 0.05; Figure S19). When considering the 30 SNPs most significantly associated with heart failure risk, 93% respond to DOX in at least one timepoint or drug concentration (adj. *P* < 0.05; Figure 6B). CX affects the expression of only three heart failure genes - *PSRC1*, *CDKN1A*, and *LSM3.* Together, these results suggest that CX does not broadly impact genes in risk loci for DOX-induced cardiotoxicity and heart failure.

## Discussion

CX is an anti-cancer drug currently in clinical trials for the treatment of advanced solid and hematological malignancies. CX has been described as an RNA polymerase I inhibitor, which takes advantage of the high levels of ribosomal RNA transcription in cancer cells. Recent studies have identified TOP2B, a topoisomerase isoform that is highly expressed in the heart, as the primary target of CX (12, 14). TOP2B is the target of several anti-cancer drugs including etoposide, mitoxantrone, DOX and related anthracyclines. DOX and mitoxantrone are known to cause adverse effects on the heart (52, 53). It has therefore been hypothesized that CX might similarly adversely affect the heart (14). We therefore sought to compare the cellular and molecular effects of CX and DOX treatment on iPSC-CMs. Our results indicate that despite similarity in drug target, CX and DOX differ in the extent of their effects on cardiomyocytes.

### CX causes less DNA damage and cytotoxicity than DOX

We first tested the effects of a range of concentrations of 48 hours of DOX and CX treatment on cardiomyocyte viability. We found that CX is approximately 20-fold less toxic to iPSC-CMs than DOX (median CX LD50 = 10.03 µM). CX is an effective anti-cancer agent that decreases cancer cell viability. EC50 values following CX treatment in ten cancer cell lines from distinct organ sites range from 0.63 - 4.7 µM suggesting that CX is less toxic to cardiomyocytes than rapidly dividing cancer cells (54, 55). The effect of DOX on iPSC-CM viability is similar to the TOP2 inhibitor anthracyclines epirubicin and daunorubicin, while the related anthracenedione, mitoxantrone, is more cytotoxic (22). These results suggest that while different drugs may similarly inhibit TOP2B, the mediator of DOX-induced cardiotoxicity, the effects on cardiomyocyte viability may differ across drugs.

At a molecular level, DOX is known to initiate the DNA damage response by inducing DNA double-strand breaks. As expected, we observe the presence of the DNA double-strand break marker, phosphorylated γ-H2AX, in most cardiomyocytes treated with 0.5 µM DOX within three hours of treatment. The proportion of cardiomyocytes with phosphorylated γ-H2AX following CX treatment is similar to VEH treatment at all concentrations tested (0.1 to 10 µM). These results demonstrate that CX does not cause DNA double-strand breaks in cardiomyocytes as DOX does, suggesting differing mechanisms of action.

### CX induces hundreds of gene expression changes that are reflected in the DOX response

Transcriptome profiling following CX treatment at two concentrations and three timepoints identified two DEGs following three hours of treatment and hundreds following 24 and 48 hours of treatment across 0.1 and 0.5 µM drug concentrations. DEGs are enriched for processes related to chromosome segregation and DNA replication at both concentrations. These same processes are enriched in iPSC-CMs treated with 0.1 µM DOX for 24 and 48 hours but not 0.5 µM DOX treatments. Joint modeling of drug responses over time identified gene expression signatures shared between CX and DOX, and specific to DOX. CX does not associate with a specific gene expression response signature. Of the 1,996 genes that respond to 0.1 µM DOX in at least one timepoint, 415 (21%) also respond to CX. However, of the 6,992 genes that respond to 0.5 µM DOX, only 363 (5%) also respond to CX. This suggests a high degree of similarity in drug responses at low concentrations but divergence at higher concentrations. When considering other TOP2 inhibitors, DOX, epirubicin, daunorubicin and mitoxantrone display strong similarity in gene expression profiles following three hours of 0.5 µM treatment, but response to the most distantly-related drug, mitoxantrone, diverges from the anthracyclines following 24 hours of treatment (22).

Divergence in CX and DOX expression profiles may result from differential effects on transcriptional regulators. DOX treatment induces the expression of the DNA damage response factor p53, and its targets, unlike CX, which implicates p53 as an upstream mediator of DOX-induced cardiotoxicity.

### TOP2B in drug-induced cardiotoxicity

Intriguingly, while both DOX and CX affect TOP2B activity, *TOP2B* mRNA levels are decreased following 24 hours of DOX, but not CX treatment, suggesting differing effects of TOP2 inhibition across drugs. TOP2B is a multifunctional protein implicated not only in DNA double-strand breaks, but also in processes including chromatin organization (56), and transcription (57). Depending on the way a drug affects TOP2B, the effects on TOP2B-related processes in the cell may differ.

Inhibition of TOP2B in both *in vivo* and *in vitro* models has indicated that TOP2B mediates DOX-induced cardiotoxicity (18, 21). The *RARG* gene, which is in a locus associated with DOX-induced cardiotoxicity, is affected by the expression of TOP2B, and RARG agonists improve DOX-induced cardiotoxicity (58). In our experiments *RARG* expression is increased following 24 and 48 hours of 0.5 µM DOX treatment but not CX treatment. TOP2B expression levels are decreased in PBMCs within hours of DOX treatment in human patients further suggesting an association with the treatment (59). TOP2B is also the target of Dexrazoxane, currently the only FDA-approved drug used to mitigate the effects of DOX on the heart. It has been shown to degrade TOP2B and therefore can be administered prior to DOX treatment to decrease cardiotoxicity (60). However, the use of Dexrazoxane is limited because it reduces anthracycline anti-tumor activity (61). This could relate to the fact that Dexrazoxane can affect both TOP2A and TOP2B isoforms (62). Dexrazoxane does not improve DOX-induced cardiotoxicity in iPSC-CMs suggesting further unknown aspects of DOX-induced cardiotoxicity (20). Indeed, there are competing hypotheses for the mechanism behind DOX-induced cardiotoxicity including ferroptosis (63). Our results provide further support for the notion that DOX induces cardiotoxicity through several mechanisms and not TOP2B alone.

### Limitations of the study

iPSC-CMs tend to resemble fetal, rather than adult cardiomyocytes. We use 30-day-old cardiomyocytes that are cultured in glucose-free media to mitigate these effects. Nevertheless, it is possible that the effects of CX treatment may differ between iPSC-CMs and adult human cardiomyocytes. For example, our iPSC-CMs express both the *TOP2A* and *TOP2B* isoforms, while human adult left ventricle and right atrium heart tissue predominantly express the *TOP2B* isoform (44, 64).

We selected two sub-micromolar drug concentrations and three treatment times to study effects on the transcriptome. Different effects may be observed at different drug concentrations or times. Indeed, we observe a stronger overlap between CX and DOX responses at 0.1 µM concentrations compared to 0.5 µM. While DOX is used in cancer patients at a dose within the range of our *in vitro* concentrations, CX is still under investigation as a potential anti-cancer drug; however the concentrations we used were in line with the plasma concentrations observed in clinical trial patients. We did not measure long-term effects of these drugs and it therefore could be the case that CX induces cardiotoxicity cumulatively over time.

We measured the effects of CX on cell viability, DNA damage and the transcriptome. CX may affect other molecular phenotypes including protein expression and activity, or cellular phenotypes including metabolism and contraction. However, the reported roles of CX in RNA polymerase I transcription inhibition, TOP2 inhibition and G-quadruplex stability would suggest that its impacts would be detected at the transcriptional level.

## Conclusions

In summary, we characterized the human cardiomyocyte response to CX treatment and contrasted it to the well-known effects of DOX. We found CX to be an order of magnitude less cytotoxic and DNA damage-inducing than DOX. These effects are evident on the transcriptome where CX induced minimal changes in gene expression and did not affect the expression of genes in loci associated with DOX-induced cardiotoxicity. Our results indicate that CX does not adversely affect the heart, suggesting that it can be further considered for use in human cancer patients.

### Code availability

All custom analysis scripts used for this project are available at https://github.com/mward-lab/Paul_CX_2025.

## Acknowledgements

We thank all members of the Ward Lab for helpful discussions. We thank Kelly Frazer and the University of California San Diego for providing the iPSC lines through the iPSCORE resource. We thank the Next Generation Sequencing Core Facility at the University of Texas Medical Branch for preparing and sequencing the RNA-seq libraries, and the Flow Cytometry Core Facility for access to flow cytometers. The authors acknowledge the Texas Advanced Computing Center (TACC) at The University of Texas at Austin for providing HPC resources that have contributed to the research results reported within this paper (http://www.tacc.utexas.edu). The Genotype-Tissue Expression (GTEx) Project was supported by the Common Fund of the Office of the Director of the National Institutes of Health, and by NCI, NHGRI, NHLBI, NIDA, NIMH, and NINDS. The data used for the analyses described in this manuscript were obtained from the GTEx Portal on 05-06-2025. This work was supported by a CPRIT Scholar award to M.C.W.

## Author contributions

M.C.W conceived and designed the study. S.P., J.A.G., A.B. performed experiments. S.P. and M.C.W analyzed the data. S.P. and M.C.W wrote the manuscript with input from co-authors. M.C.W supervised the work.

## Ethics approval

The six iPSC lines were generated with approval from the Institutional Review Boards of the University of California, San Diego and The Salk Institute (Project no. 110776ZF) and informed written consent of participants. Cell lines are available through the biorepository at WiCell Research Institute (Madison, WI, USA), or through contacting Dr. Kelly A. Frazer at the University of California, San Diego.

## Supplemental material

S1 Appendix: Document containing Supplemental Figures 1-19

S2 Appendix: Document containing Supplemental Tables 1-16

