## Supplementary material for "Topoisomerase II inhibitors CX-5461 and Doxorubicin differ in their cardiotoxicity profiles": S1 Appendix

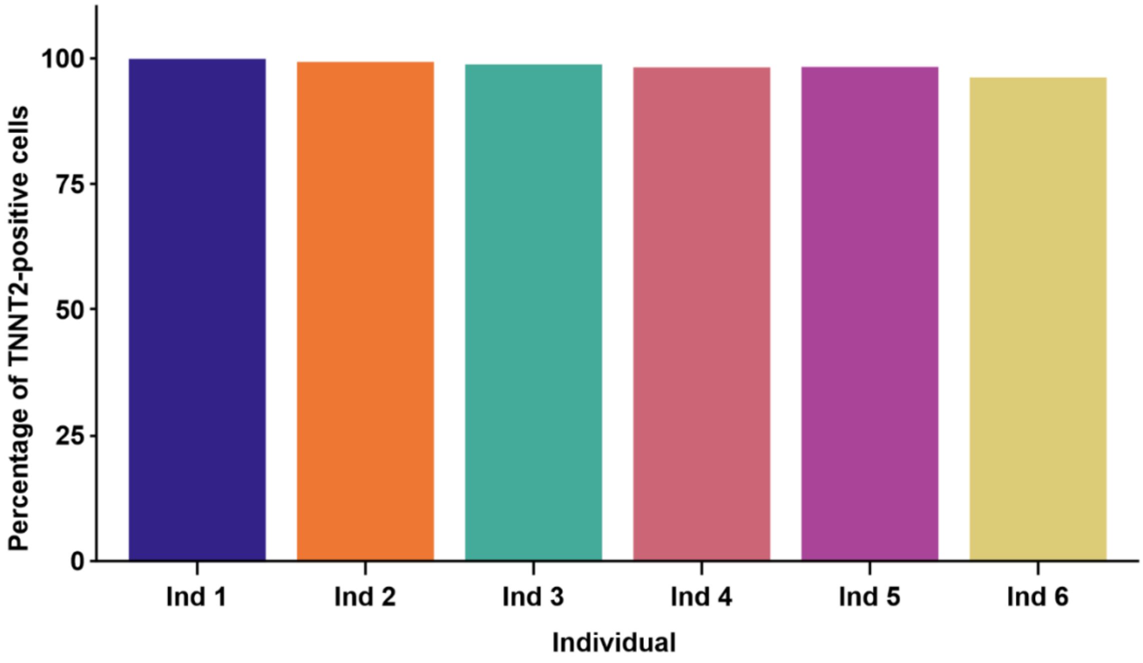


**Figure S1: iPSCs can be successfully differentiated into cardiomyocytes.** Proportion of cells in each individual expressing cardiac Troponin T as measured by flow cytometry.

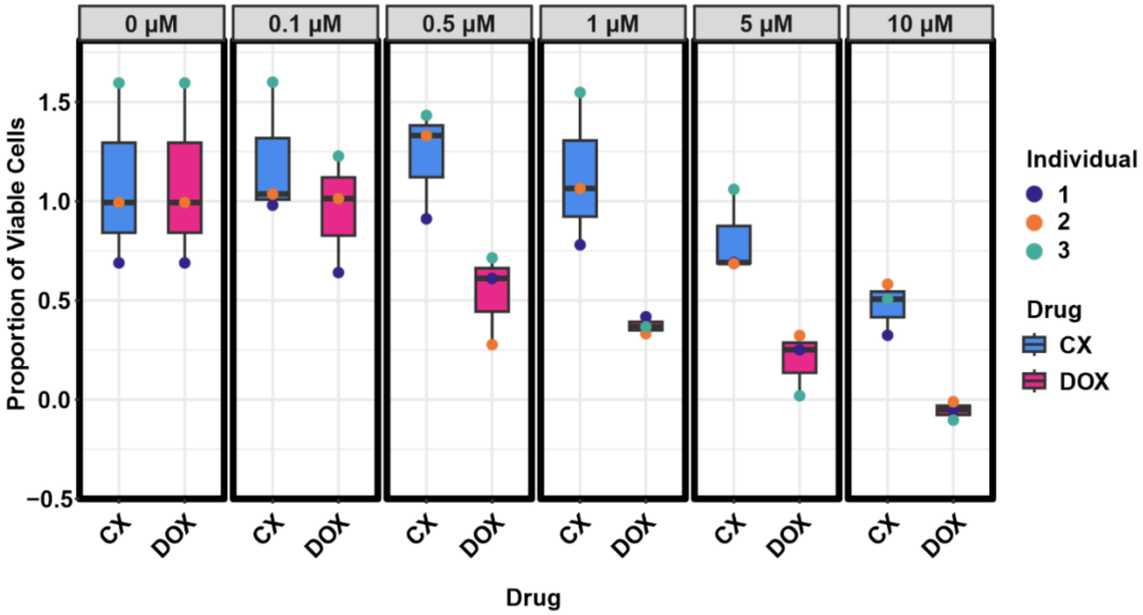


**Figure S2: CX is less cytotoxic than DOX.** Proportion of viable cells following treatment with increasing concentrations of CX and DOX for 48 hours. Values represent viability of the drug-treated cells at each timepoint relative to VEH. Measurements are from Individuals 1,2 and 3.

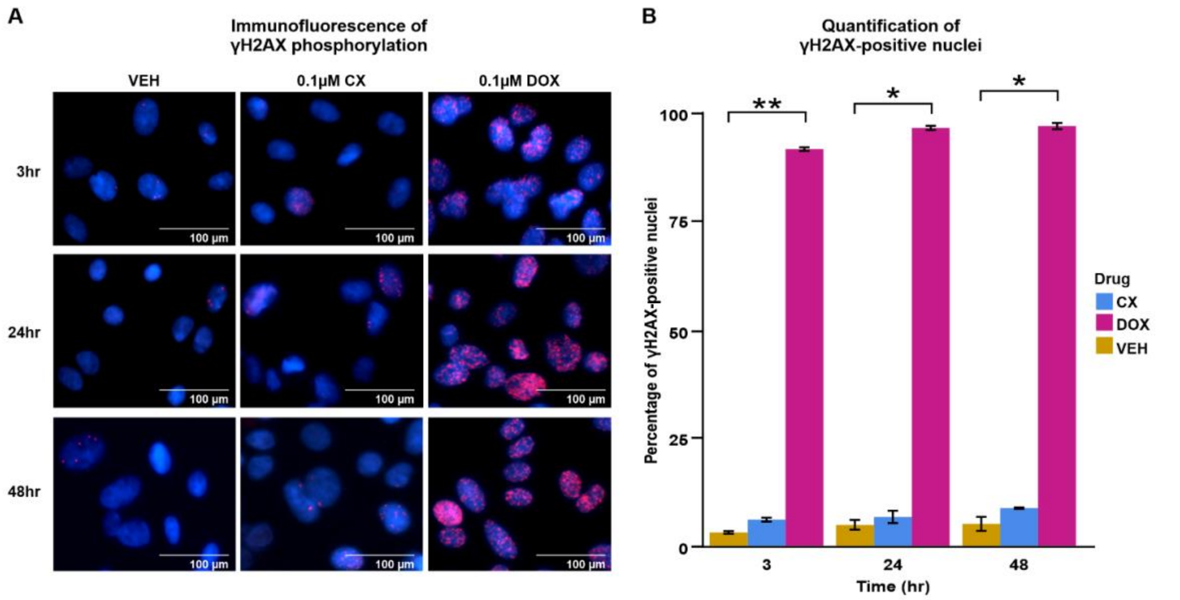


**Figure S3: Low dose DOX treatment induces DNA damage unlike CX. (A)** γH2AX (red) and Hoechst DNA stain (blue) in iPSC-CMs treated with VEH, 0.1 μM CX or 0.1 μM DOX for three, 24 or 48 hours. Scale bar: 100 μm. **(B)** Proportion of iPSC-CMs that are positive for γH2AX. Data representative of 100 cells per treatment per individual. Data are presented as mean ± SD across two individuals (Individuals 2 and 3). Asterisk represents a statistically significant difference in γH2AX expression (**P* < 0.05, ***P* < 0.01).

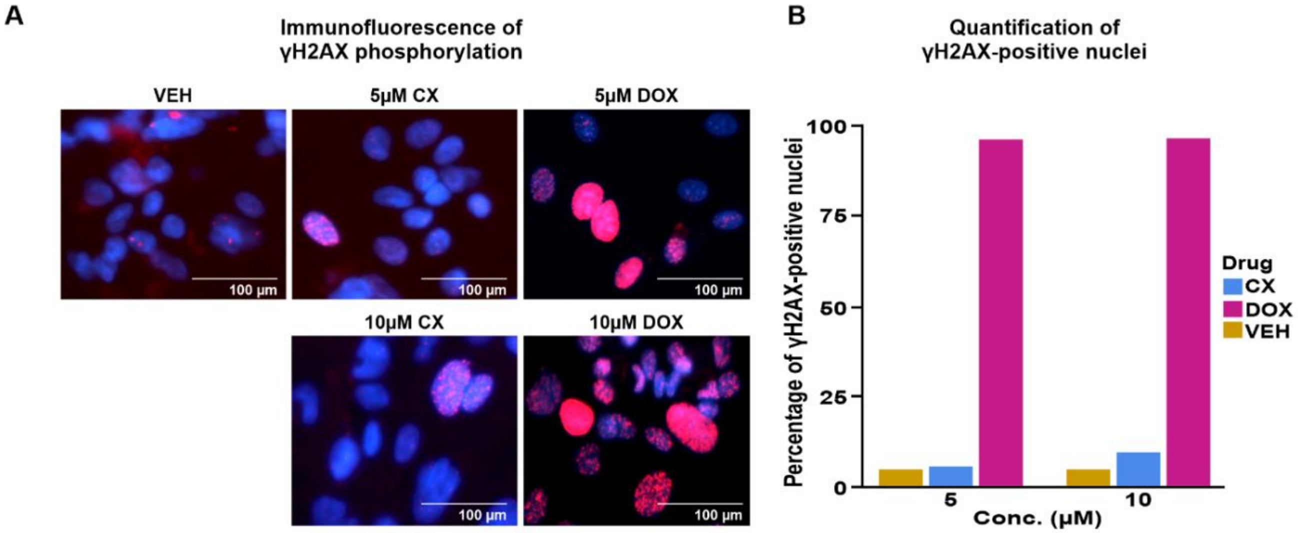


**Figure S4: High dose CX treatment induces minimal DNA damage. (A)** γH2AX (red) and Hoechst DNA stain (blue) in iPSC-CMs treated with 0, 5 or 10 μM CX or DOX for 24 hours. Scale bar: 100 μm. **(B)** Proportion of cells positive for γH2AX. Data representative of 100 cells per treatment from one individual (Individual 3).

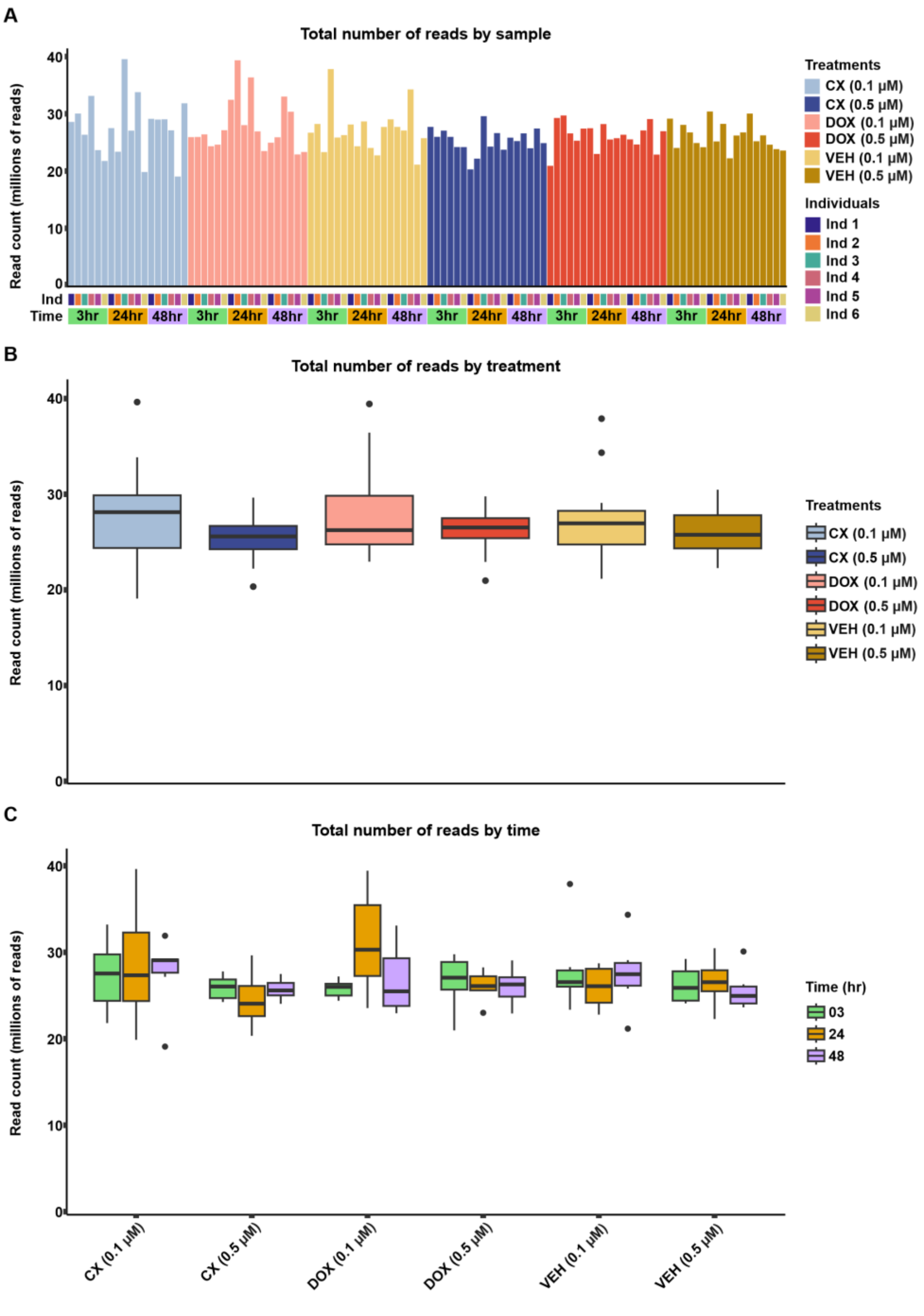


**Figure S5**: **RNA-seq read number is similar across treatment groups and time. (A)** Total number of RNA-seq read pairs for each sample. **(B)** Total number of RNA-seq read pairs per treatment group. **(C)** Total number of RNA-seq read pairs per treatment and time group.

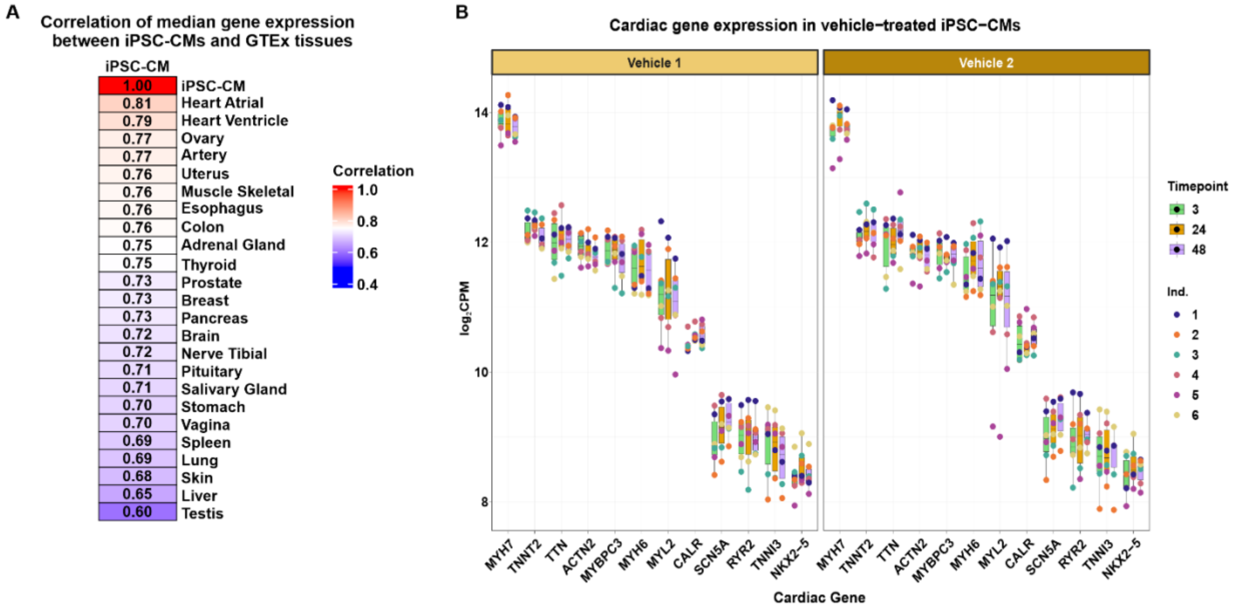


**Figure S6**: **iPSC-CM gene expression profile resembles human heart tissue. (A)** Pearson correlation between median expression of VEH-treated iPSC-CMs, and expression across 24 human tissues (GTEx Consortium, 2020). **(B)** Expression of a panel of heart marker genes across VEH-treated iPSC-CMs. Vehicle 1 and Vehicle 2 represent DMSO-treated control samples volume-matched to the 0.1 µM and 0.5 µM drug treatments respectively.

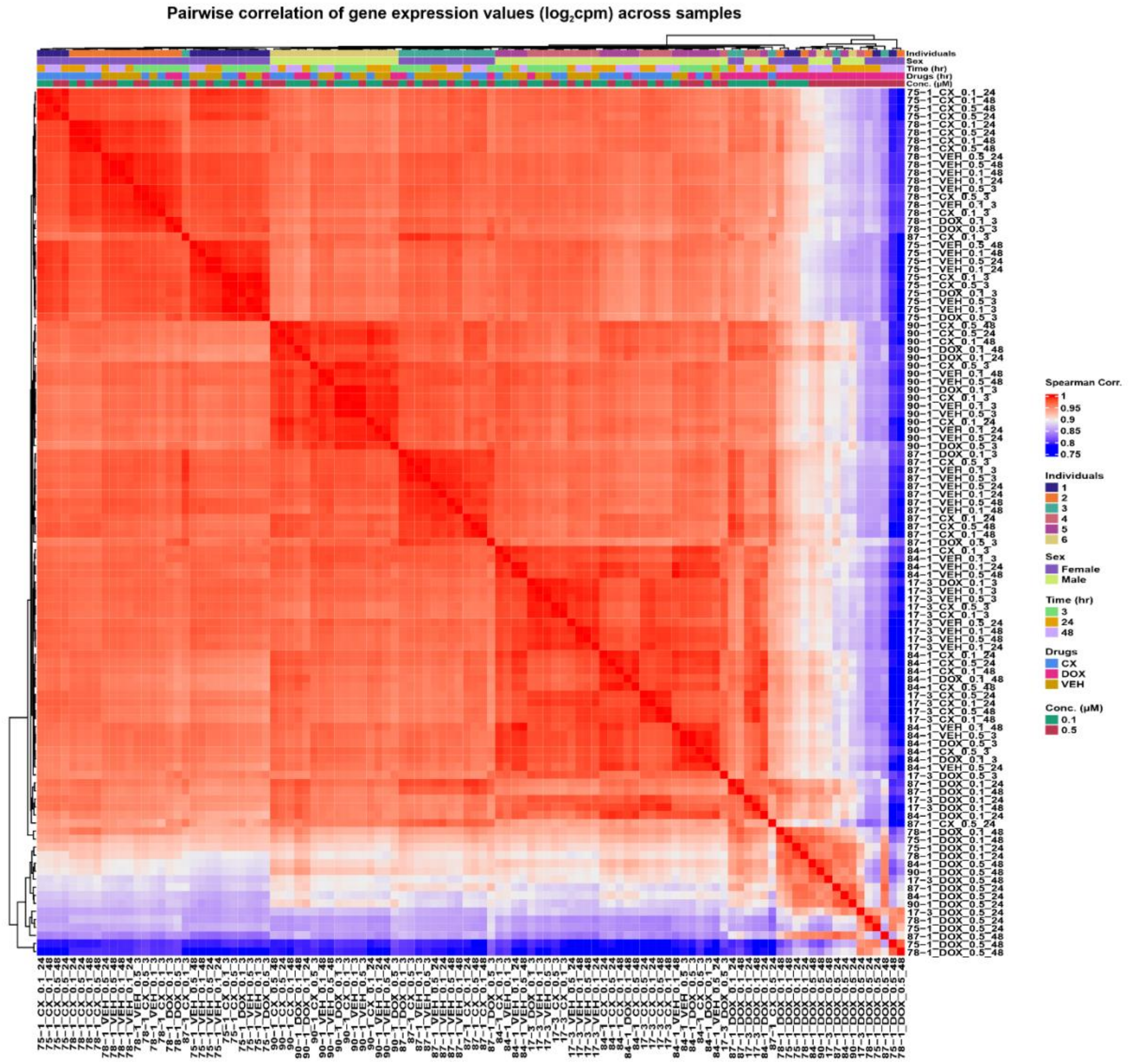


**Figure S7**: **RNA-seq samples cluster by treatment type, drug concentration, timepoint, and individual.** Spearman correlation of $\log_2$cpm values across all pairs of samples.

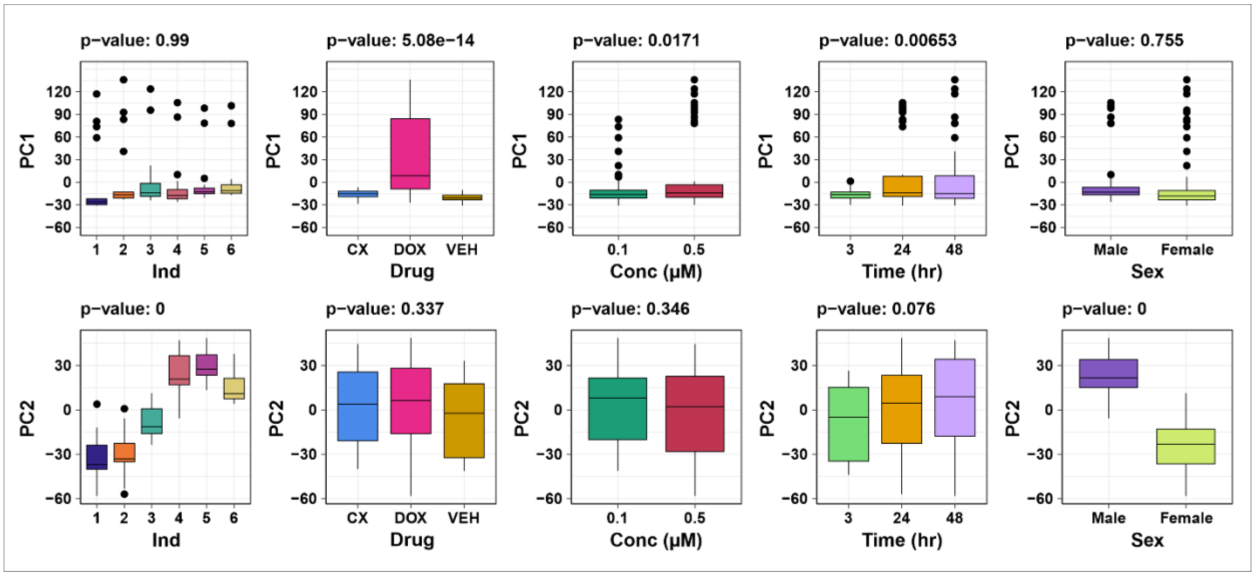


**Figure S8**: **PC1 associates with treatment, drug concentration and treatment time, while PC2 associates with individual and sex.** Demonstration of variance contributed to the first two principal components from five major covariates in the study: individual, treatment, drug concentration, treatment time and sex. The correlation between each covariate and each PC is calculated using a linear model. *P* values represent the significance of the F-statistic from the model.

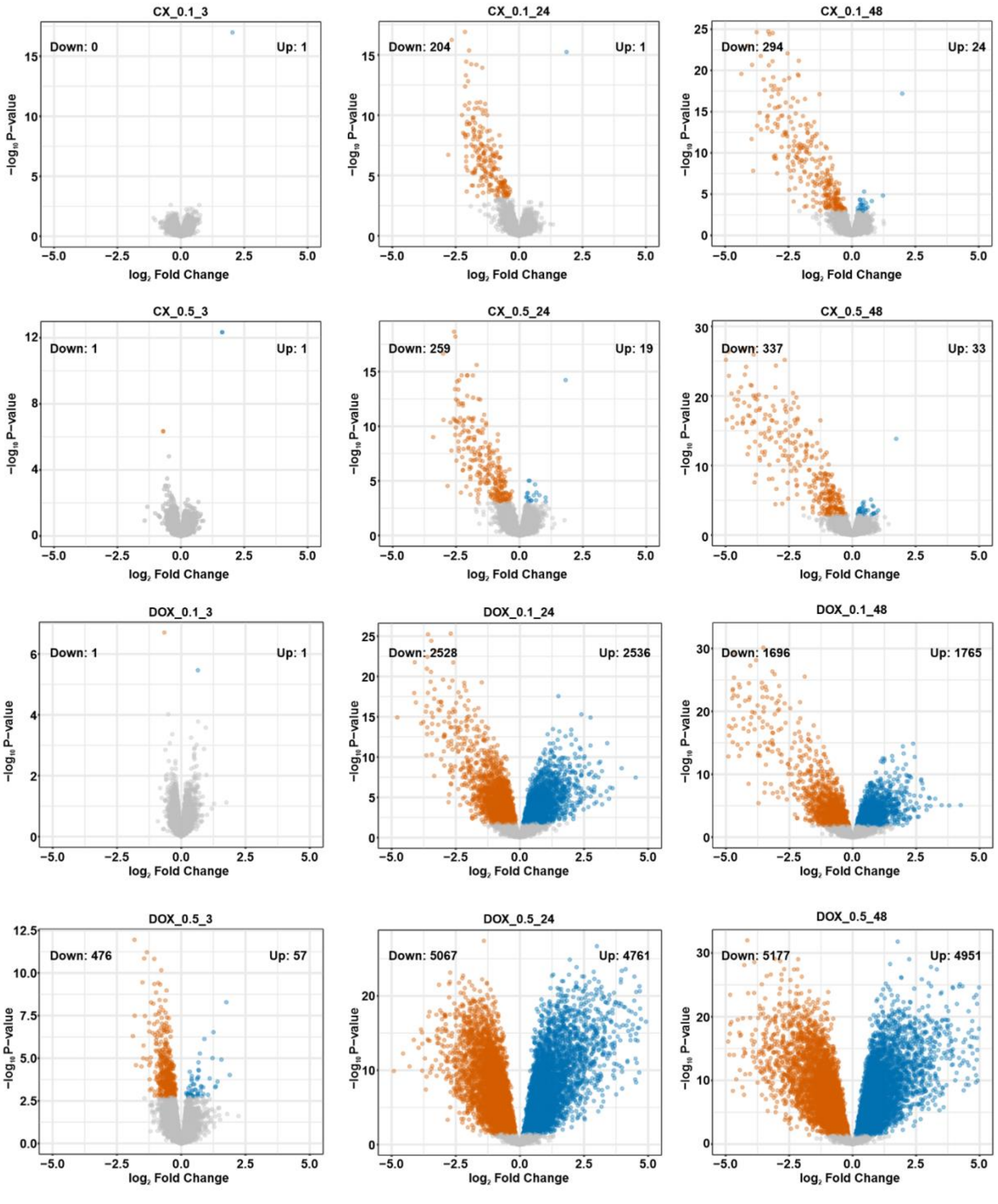


**Figure S9**: **Hundreds of gene expression changes are induced in response to CX treatment over time.** Volcano plots representing genes that are differentially expressed between CX or DOX (0.1 or 0.5 μM) and VEH treatment at each timepoint (three, 24 or 48 hours). Genes that are significantly up-regulated in response to treatment (adj. $P < 0.05$) are represented in blue, and genes that are significantly down-regulated are represented in vermilion. The number of genes that are up- and down-regulated is given for each plot.

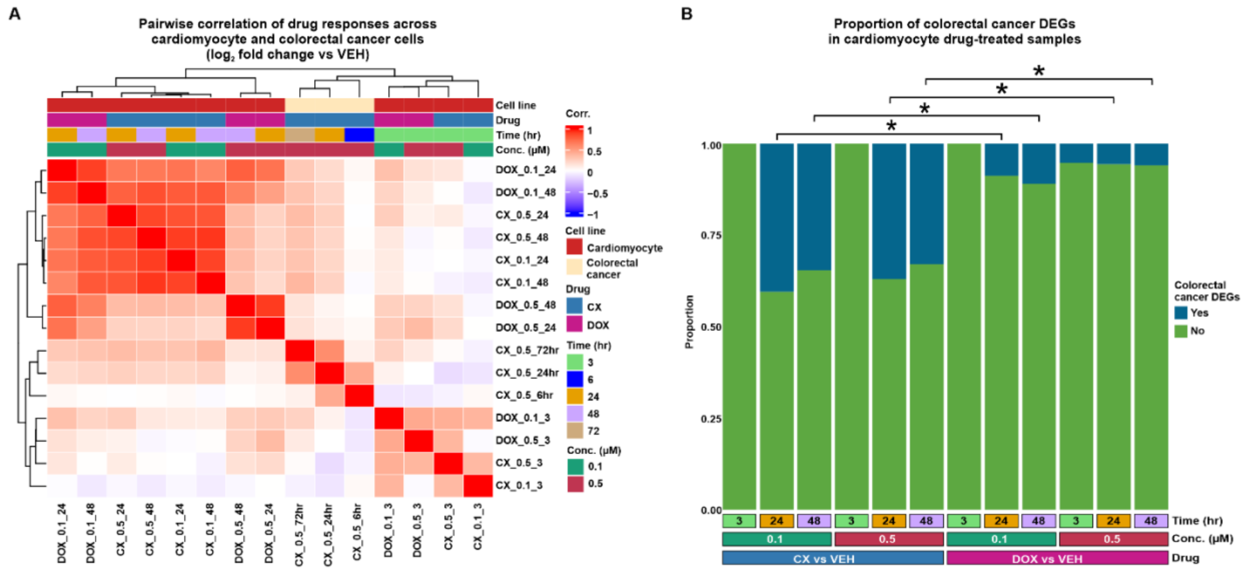


**Figure S10**: **Gene expression response to CX treatment in iPSC-CMs resembles the response in colorectal cancer cells. (A)** Correlation of drug response effect sizes ($log_2$ fold change drug vs VEH) for our data in iPSC-CMs and colorectal cancer cells treated with 0.5 µM CX for six, 24 or 72 hours (Otto *et al*., 2022). **(B)** Proportion of CX and DOX DEGs that are CX DEGs in colorectal cancer cells. Asterisk represents a significant difference in the proportion of colorectal cancer CX DEGs between iPSC-CM CX and DOX DEGs in each treatment.

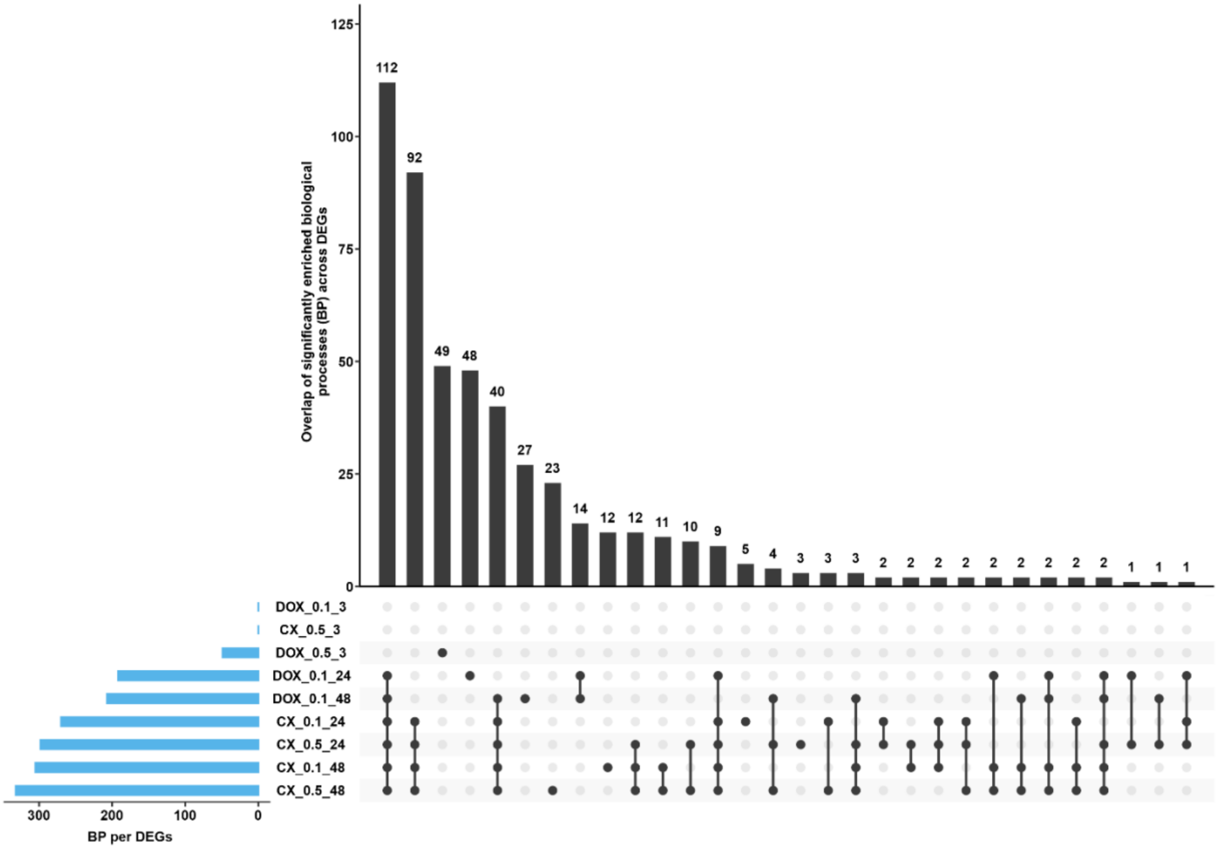


**Figure S11**: **Biological processes enriched in CX and DOX DEGs are often shared across treatments.** Intersection between all enriched biological processes (adj. $P < 0.05$) across treatments.

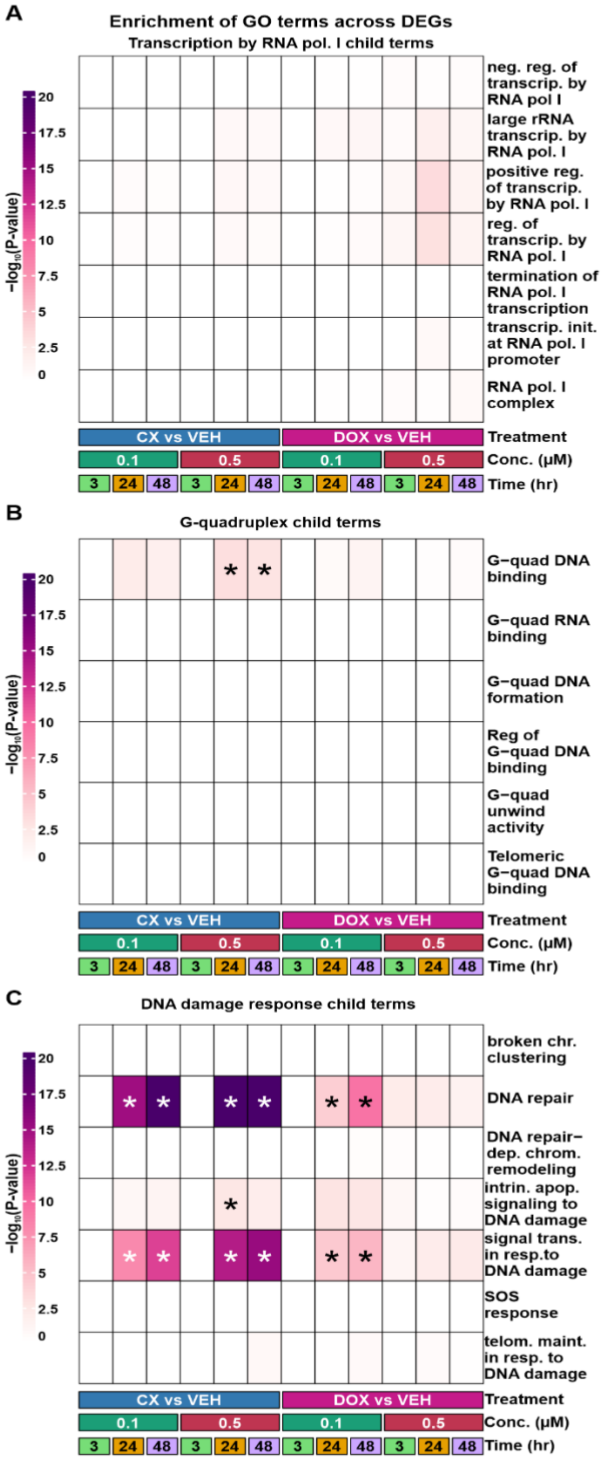


**Figure S12: G-quadruplex- and DNA damage-associated terms are enriched amongst CX response genes but not terms associated with RNA polymerase I transcription. (A)** Enrichment of all biological processes associated with 'RNA polymerase I transcription' amongst DEGs for each treatment. **(B)** Enrichment of all molecular function terms associated with 'G-quadruplex' amongst DEGs. Asterisk represents significantly enriched processes (adj. *P* < 0.05). **(C)** Enrichment of all biological processes associated with the 'DNA damage response' amongst DEGs. Asterisk represents significantly enriched processes (adj. *P* < 0.05).

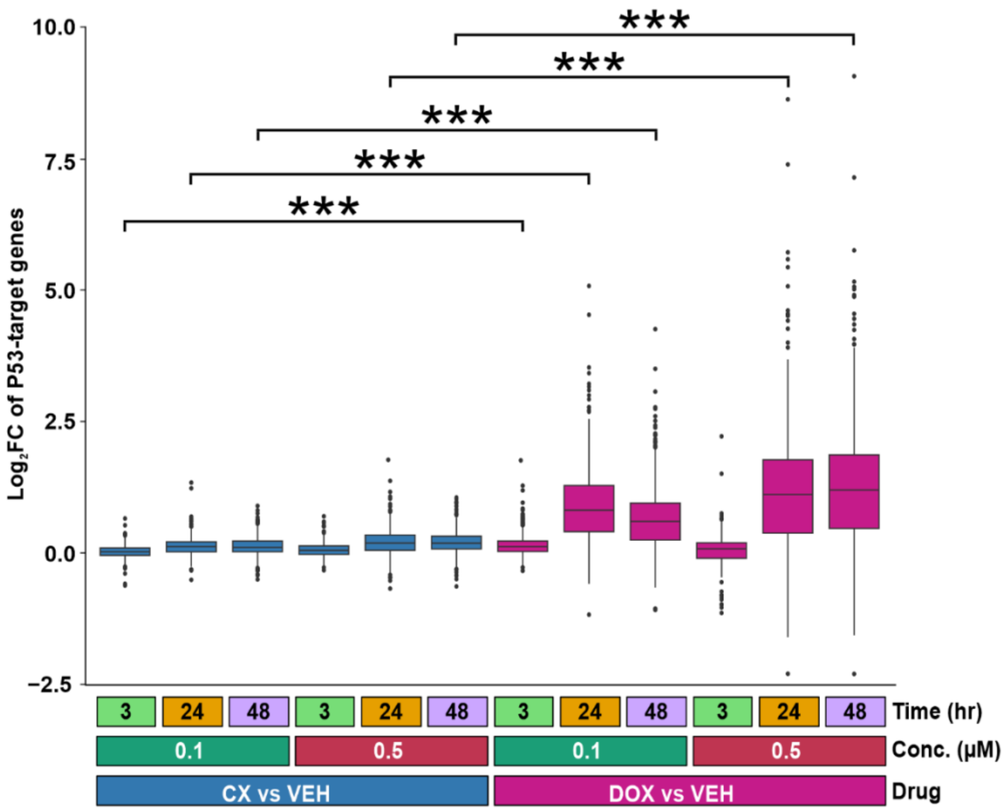


**Figure S13: DOX but not CX induces a response amongst known p53 response genes.** Drug response ($\log_2$ fold change) between each drug treatment and VEH of a set of 300 p53 target genes (Fischer *et al*., 2017). Asterisk represents a significant difference between CX and DOX responses (****P* < 0.001).

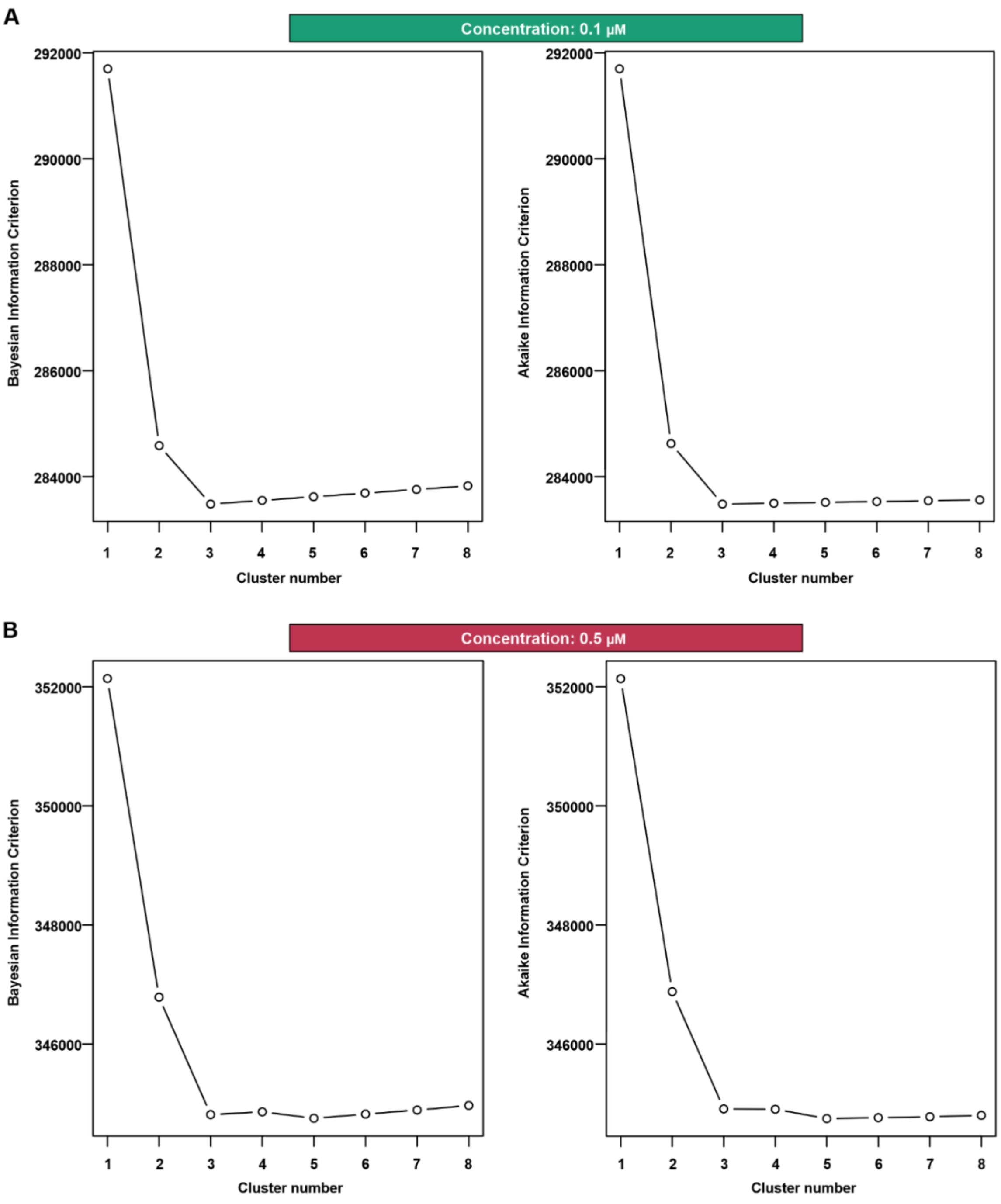


**Figure S14: Three gene expression clusters capture the response to 0.1 µM drug treatment and five capture the response to 0.5 µM drug treatment. (A)** Bayesian information criterion (BIC) and Akaike information criterion (AIC) at increasing numbers of Cormotif correlation motifs following joint modeling of pairs of tests for the 0.1 µM drug treatments. **(B)** BIC and AIC at increasing numbers of Cormotif correlation motifs following joint modeling of pairs of tests for the 0.5 µM drug treatments.

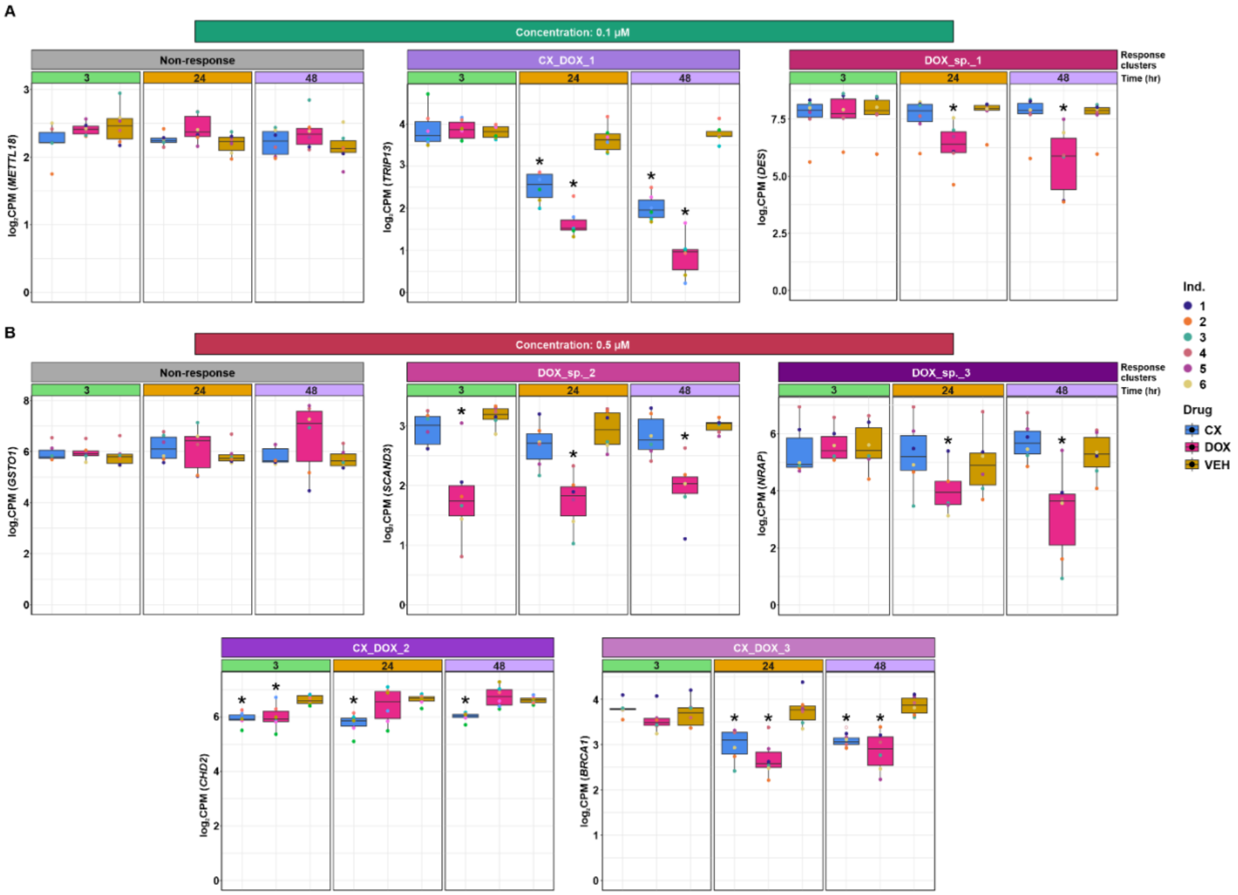


**Figure S15: Gene expression response clusters identify genes that respond to treatment. (A)** Gene expression levels of genes assigned to each of the three drug response clusters for each 0.1 µM drug treatment at each time point. **(B)** Gene expression levels of genes assigned to each of the five drug response clusters for each 0.5 µM drug treatment at each time point. Asterisk represents treatments where the gene is classified as a DEG (adj. $P < 0.05$).

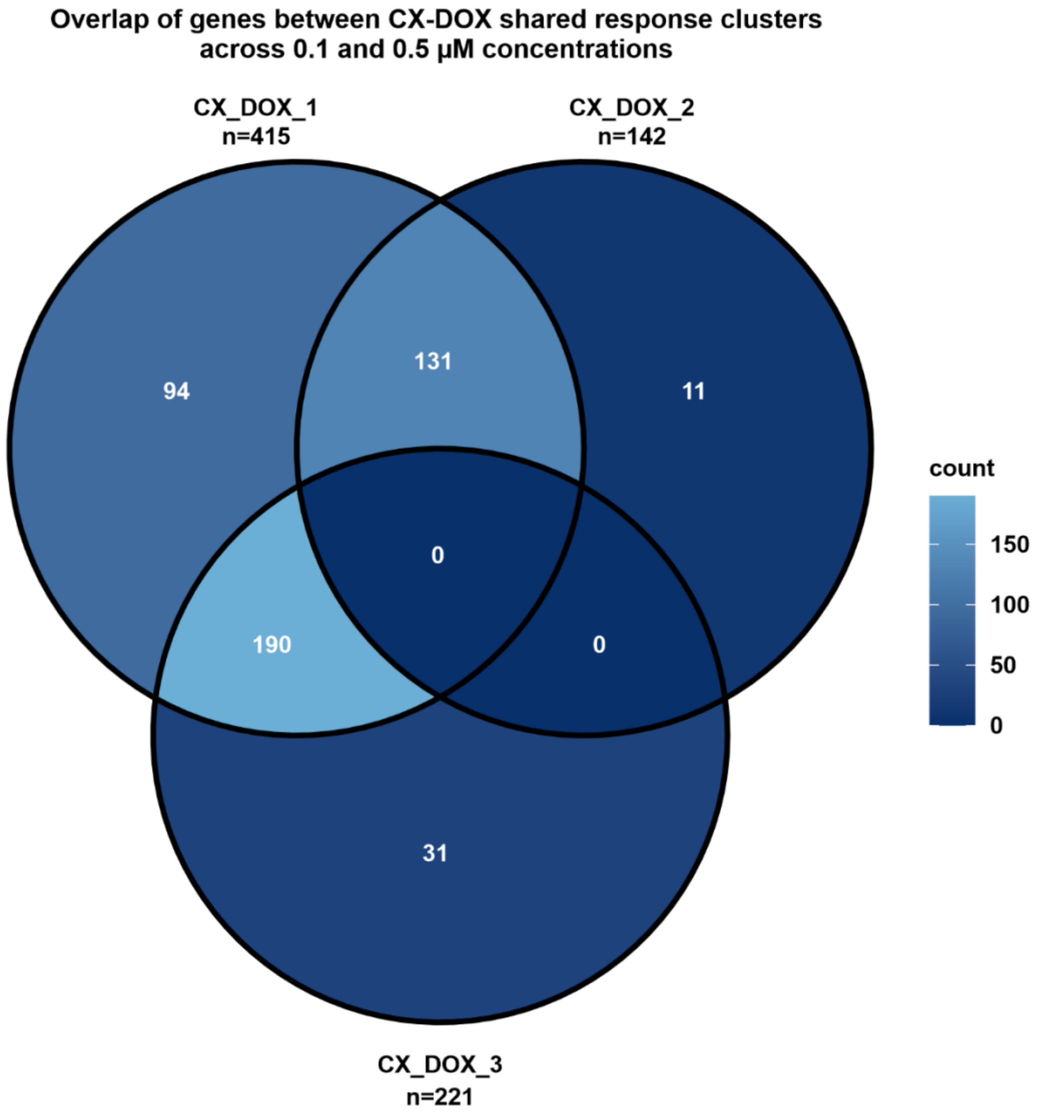


**Figure S16: Most genes that respond to 0.1 μM CX treatment also respond to 0.5 μM CX treatment.** Overlap between the three CX response clusters across drug concentrations (CX_DOX_1: 0.1 μM, CX_DOX_2: 0.5 μM, CX_DOX_3: 0.5 μM).

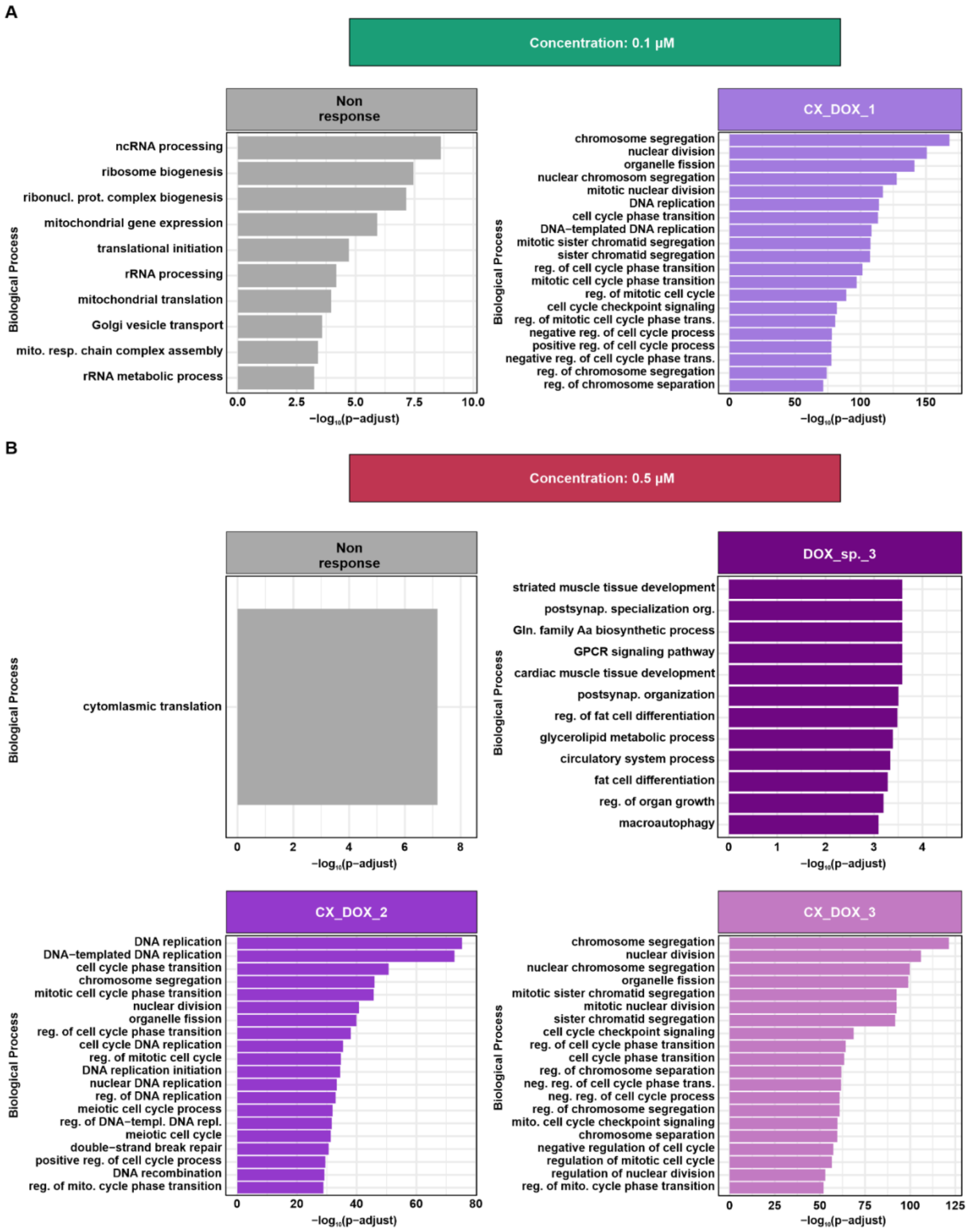


**Figure S17: CX response clusters enrich for biological processes related to chromatin segregation and DNA replication. (A)** Top 20 enriched biological processes (adj. $P < 0.05$) for each 0.1 μM drug response cluster. DOX_sp._1 cluster has no enriched biological processes. **(B)** Top 20 enriched biological processes (adj. $P < 0.05$) for each 0.5 μM drug response cluster. DOX_sp._2 cluster has no enriched biological processes.

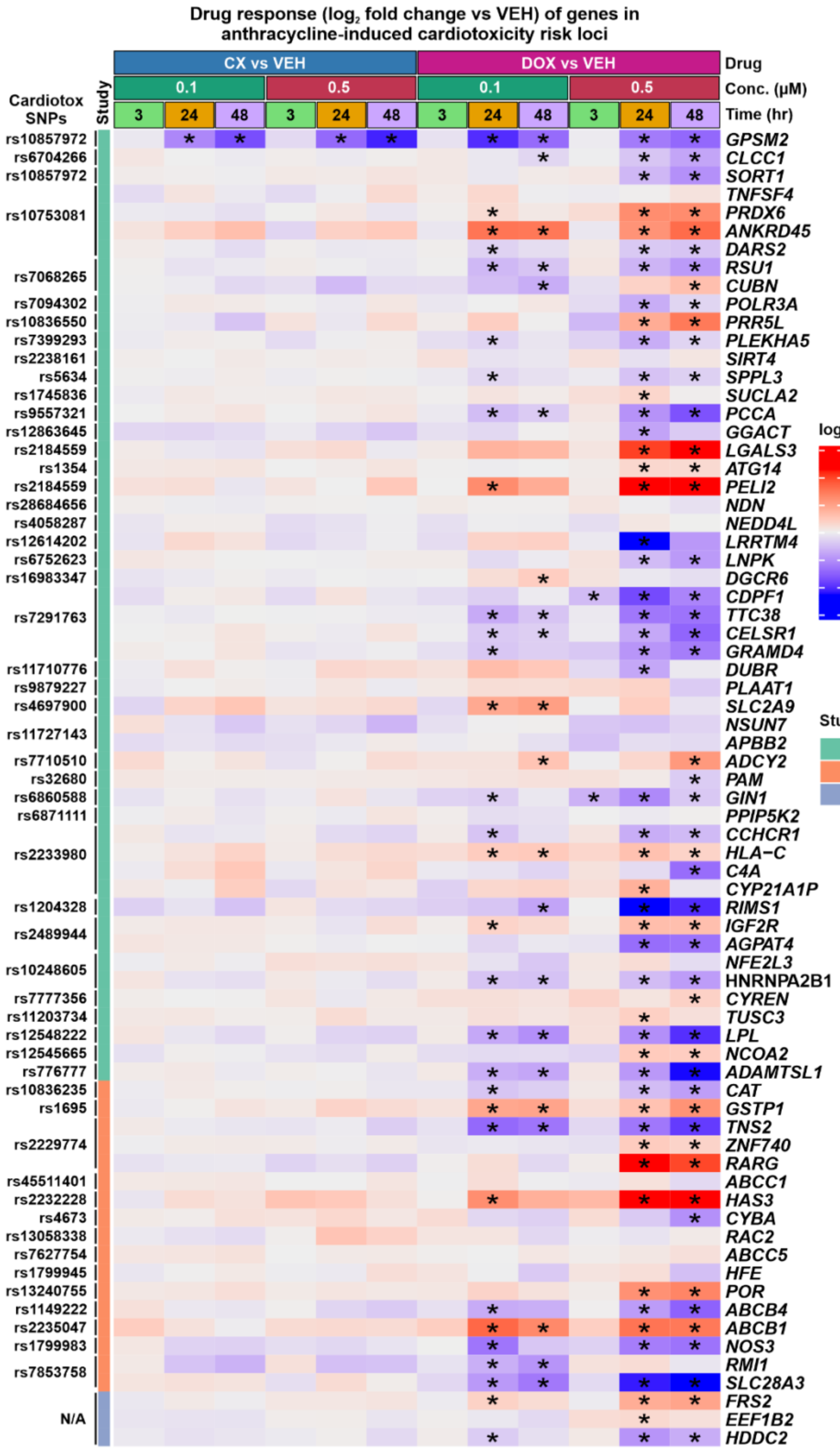


**Figure S18: Most genes in anthracycline-induced cardiotoxicity loci do not respond to CX treatment.** Drug responses ($log_2$ fold change) of genes in anthracycline-associated cardiotoxicity risk loci (Schneider *et al*., 2017: GWAS1; Aminkeng *et al*., 2016: GWAS2; Scott *et al*., 2021: TWAS). GWAS SNPs were associated with genes based on whether they are the closest gene to the SNP or are an eQTL in any tissue. Asterisk represents genes that are classified as DEGs (adj. $P < 0.05$).

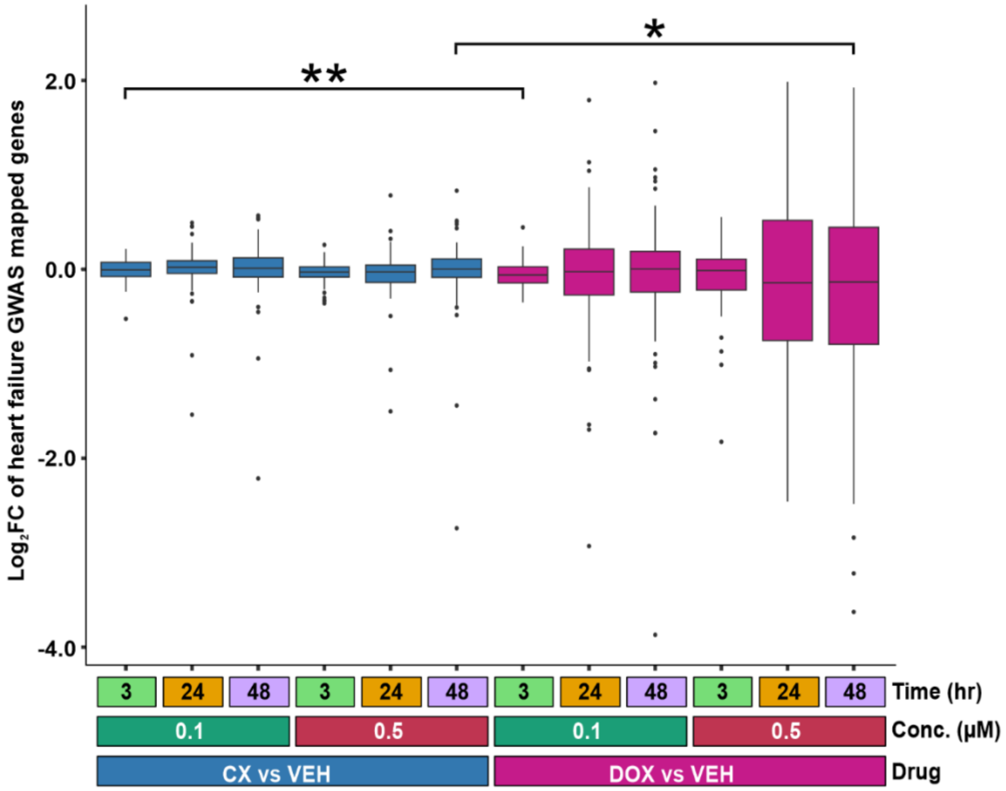


**Figure S19: Genes in loci associated with heart failure respond to DOX but not CX treatment.** Drug response ($\log_2$ fold change) between each drug treatment and VEH of mapped genes in loci associated with risk for heart failure (Cerezo *et al*., 2025). Asterisk represents a significant difference between CX and DOX responses (*$P < 0.05$ and **$P < 0.01$).
